# Extracellular electron transfer increases fermentation in lactic acid bacteria via a hybrid metabolism

**DOI:** 10.1101/2021.05.26.445846

**Authors:** Sara Tejedor-Sanz, Eric T. Stevens, Peter Finnegan, James Nelson, Andre Knoessen, Samuel H. Light, Caroline M. Ajo-Franklin, Maria L. Marco

**Affiliations:** Department of Biosciences, Rice University, Houston, United States; Biological Nanostructures Facility, The Molecular Foundry, Lawrence Berkeley National Laboratory, Berkeley, United States; Department of Food Science & Technology, University of California-Davis, Davis, United States; Department of Electrical and Computer Engineering, University of California-Davis, Davis, United States; Department of Microbiology, University of Chicago, Chicago, United States

## Abstract

Energy conservation in microorganisms is classically categorized into respiration and fermentation, however recent work shows some species can use mixed or alternative bioenergetic strategies. We explored the utility of a flavin-based extracellular electron transport (FLEET) system for energy conservation within diverse lactic acid bacteria (LAB), microorganisms that mainly rely on fermentative metabolism and are important in food fermentations. The LAB *Lactiplantibacillus plantarum* uses extracellular electron transfer to increase its NAD^+^/NADH ratio, generate more ATP through substrate-level phosphorylation and accumulate biomass more rapidly. This novel, hybrid metabolism was dependent on a type-II NADH dehydrogenase (Ndh2) and conditionally required a flavin-binding extracellular lipoprotein (PplA) in the FLEET system to confer increased fermentation yield, metabolic flux, and environmental acidification in both laboratory media and food fermentation. The discovery of a single pathway that blends features of fermentation and respiration expands our knowledge of energy conservation metabolism and provides immediate biotechnology applications.

## INTRODUCTION

The ways in which microorganisms extract energy to maintain cellular functions are directly linked to their environment, including the availability of nutrients and cooperative or antagonistic interactions with other organisms (Haruta and Kanno, 2015). Microorganisms must also maintain redox homeostasis by responding to oxidative or reductive changes in their intracellular or extracellular environment (Sporer *et al*., 2017). Ultimately, microorganisms that can effectively generate cellular energy while also managing redox requirements will maintain higher growth and survival rates, and therefore exhibit greater ecological fitness.

All organisms possess a variety of mechanisms to utilize substrates for energy generation in the form of high-energy compounds like ATP. During respiration, microorganisms rely on either oxygen (aerobic) or other substrates (anaerobic) as terminal electron acceptors for the synthesis of ATP. Some microorganisms, most notably *Shewanella* spp. and *Geobacter* spp., can anaerobically respire using electron acceptors outside the cell, such as iron (III) oxides or an electrode (Renslow et al., 2013; Richter et al., 2012). This process is called extracellular electron transfer (EET). Regardless of the identity of the electron acceptor, ATP synthesis during respiration occurs via oxidative phosphorylation (Kim and Gadd, 2019). In oxidative phosphorylation, electrons from electron carriers are transported by an electron transport chain (ETC), which creates chemical and charge gradients and creates a proton motive force (PMF) for ATP generation. In anaerobic metabolism, energy acquisition strategies also include fermentation, an ATP-generating process in which organic compounds act as both donors and acceptors of electrons (Kim and Gadd, 2019). Fermentative bacteria produce ATP via substrate-level phosphorylation, which proceeds through the transfer of electrons via redox carriers (NADH, FADH) to endogenous organic compounds in the absence of an ETC (Tymoczko et al., 2019).

Lactic acid bacteria (LAB) are a diverse group of aerotolerant, saccharolytic microorganisms in the Firmicutes phylum that produce lactic acid as the major end-product of fermentative growth. These bacteria are essential for many food fermentations, including fermented milk and meats, fruits and vegetables, and grains (Bintsis, 2018). Strains of LAB are used for industrial chemical production (Sauer et al., 2017) and as probiotics to benefit human and animal health (Vinderola et al., 2019). Although some LAB species can perform aerobic or anaerobic respiration in the presence of exogenous heme and menaquinone, even those LAB use fermentation metabolism as the primary energy conservation pathway (Brooijmans et al., 2009). Therefore, LAB growth rates and cell yields are constrained by access to electron acceptors used to maintain intracellular redox balance during substrate-level phosphorylation.

The bioenergetics of anaerobic bacteria have been tightly linked to oxidative phosphorylation for anaerobic respiration and SLP for fermentation. However, recent experimental evidence shows a concurrent use of oxidative phosphorylation and substrate-level phosphorylation. For instance, some Crabtree-positive yeasts perform respiro-fermentation to enhance ATP production (Pfeiffer and Morley, 2014). Another example is the electron bifurcating mechanism used by some fermentative microorganisms such as *Clostridium* spp. (Herrmann et al., 2008; Li et al., 2008). Through this energy conservation strategy, cells can couple exergonic to endergonic reactions and generate extra ATP through oxidative phosphorylation (Buckel and Thauer, 2013; Müller et al., 2018). Along with other examples that are not fully understood (Hau and Gralnick, 2007; Kracke et al., 2018), these observations suggest metabolisms that combine aspects of fermentation and respiration may exist.

We recently discovered that *Listeria monocytogenes,* a facultative anaerobic pathogen known to rely on respiratory metabolism, uses EET to Fe^3+^or an anode through a flavin-based extracellular electron transport (FLEET) system (Light et al., 2018). EET in this bacterium was supported by non-fermentable and fermentable substrates. Use of FLEET allowed *L. monocytogenes* to maintain intracellular redox balance via NADH oxidation. This capacity was associated with the presence of a gene locus that was identified in many Gram-positive species in the Firmicutes phylum, including LAB. Studies in individual species of LAB such as *Lactococcus lactis* (Freguia et al., 2009; Masuda et al., 2010), *Enterococcus faecalis* (Hederstedt et al., 2020; Keogh et al., 2018), and *Lactiplantibacillus pentosus* (Vilas Boas et al., 2015) show that they can perform EET with an anode. These observations are quite surprising because EET is associated with non-fermentative respiratory organisms. They also raise the question of whether the FLEET locus is functional in LAB and what, if any, role it plays in energy conservation and metabolism.

Here, we explored EET across LAB and studied the implications of this trait at a metabolic and energetic level in *Lactiplantibacillus plantarum*, a generalist LAB species found in insect, animal, and human digestive tracts and essential for the production of many fermented foods (Behera et al., 2018; Duar et al., 2017). These findings have implications for the understanding of alternative energy generation strategies in primarily fermentative microorganisms and on lactic acid fermentations in food biotechnology.

## RESULTS

### *L. plantarum* reduces extracellular electron acceptors

To determine whether *L. plantarum* can reduce extracellular electron acceptors, we first measured its ability to reduce insoluble ferrihydrite (iron (III) oxyhydroxide). Incubation of the model strain *L. plantarum* NCIMB8826 in the presence of ferrihydrite showed that this strain reduces Fe^3+^ to Fe^2+^ (**Fig 1A and Fig S1A**). Viable cells are required for iron reduction and this activity is dependent on the presence of exogenous quinone (DHNA, 1,4-dihydroxy-2-naphthoic acid) (**Fig 1A and Fig S1A**). The requirement for DHNA was hypothesized because DHNA is a precursor of demethylmenaquinone (DMK), a membrane electron shuttle utilized by *L. monocytogenes* for EET (Light et al., 2018), and *L. plantarum* lacks a complete DHNA biosynthetic pathway (Brooijmans et al., 2009). The iron reduction was not dependent on *L. plantarum* growth medium (**Fig S1B and Fig S1C**), but for full activity, an electron donor (such as mannitol or glucose) was required to be present (**Fig 1A and Fig S1A**). Like *L. monocytogenes* (Light et al., 2018), the addition of riboflavin also increased Fe^3+^ reduction in a dose-dependent manner (**Fig S1D**). Thus, *L. plantarum* reduces insoluble iron in a manner similar to *L. monocytogenes*.

**Figure 1.**
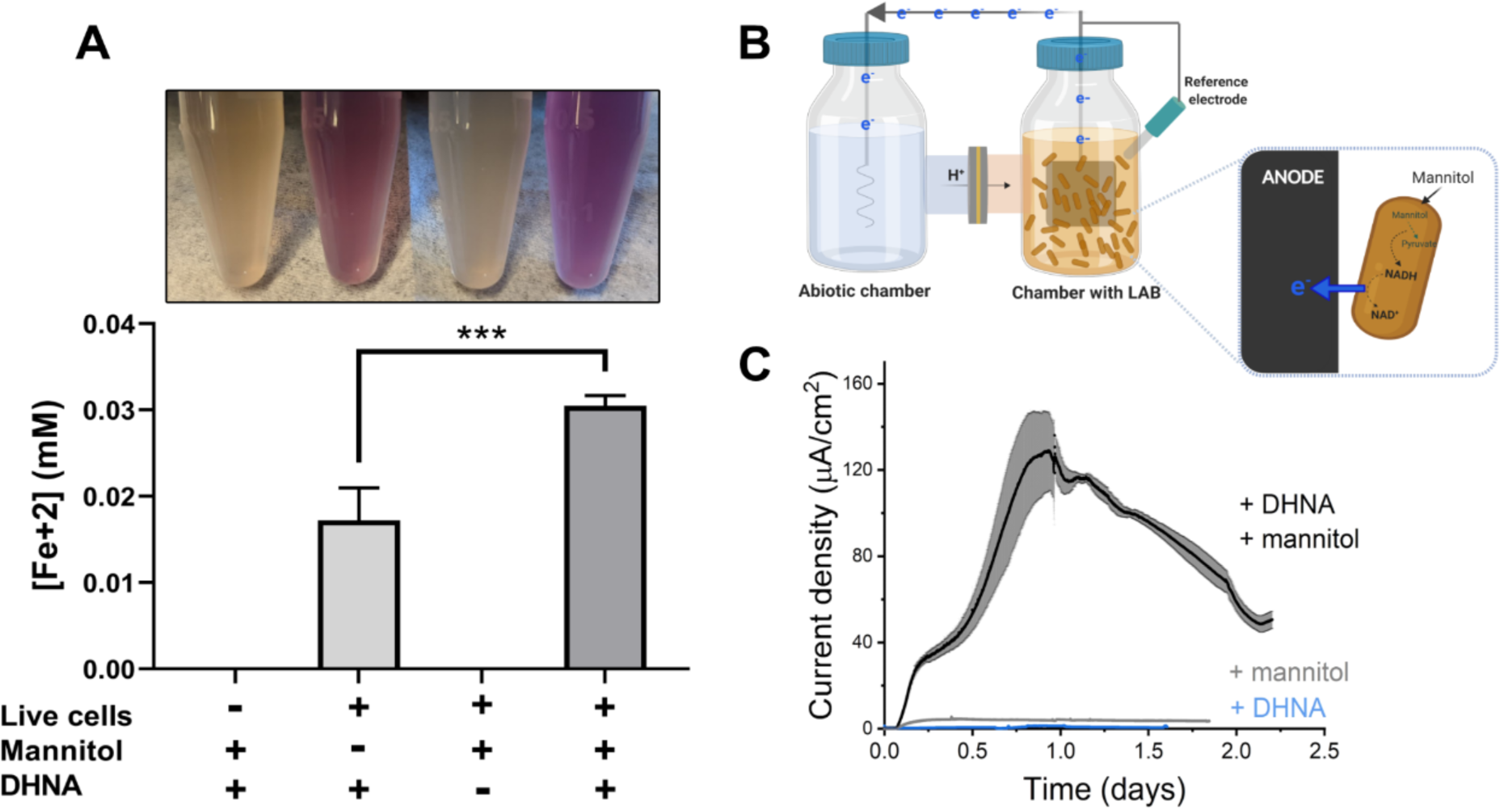
*L. plantarum* can reduce both Fe^3+^ and an anode through EET. **(A)** Reduction of Fe^3+^ (ferrihydrite) to Fe^2+^ by *L. plantarum* NCIMB8826. The assays were performed in PBS and 20 µg/mL DHNA and 55 mM mannitol were added as indicated. Fe^2+^ was detected colorimetrically using 2 mM ferrozine. For *L. plantarum* inactivation, cells were incubated at 85 ℃ in PBS for 30 min prior to the assay. Significant differences were determined by one-way ANOVA with Tukey’s post-hoc test (n = 3), *** p ≤ 0.001. **(B)** Two-chambered electrochemical cell setup for measuring current generated by *L. plantarum* through FLEET. **(C)** Current density production over time by *L. plantarum* in CDM with supplementation of 110 mM mannitol and/or 20 µg/mL DHNA. The anode was polarized at +0.2V_Ag/AgCl_. The avg ± stdev of three replicates is shown.

*L. plantarum* EET activity was confirmed in a bioelectrochemical reactor by quantifying electron output as current (**Fig 1B**). *L. plantarum* reduced a carbon electrode (anode) polarized to +200 mV_Ag/AgCl_ in the presence of both DHNA and an electron donor (mannitol) (**Fig 1C**). No current was observed in the absence of *L. plantarum* (**Fig S2A**), indicating that current production stems from a biological process. *L. plantarum* produced a maximum current of 129 ± 19 µA/cm^2^ in chemically defined medium (CDM) (**Fig 1C**), and 167 ± 26 µA/cm^2^ in de Mann Rogosa Sharpe (MRS) complete medium (De MAN et al., 1960) (**Fig S2B**). Current density was 6- to 8-fold higher than current previously observed for *L. monocytogenes* (Light et al., 2018). Remarkably, the current density capacity of *L. plantarum* is comparable to that described for *Geobacter sulfurreducens*, the model species for direct EET (140-240 µA/cm^2^) (Dumas et al., 2007; Yi et al., 2009) and significantly higher than that of *Shewanella oneidensis*, the model species for mediated EET (32-60 µA/cm^2^) (Kouzuma et al., 2014; Marsili et al., 2008). Similar to our iron reduction experiments, current production was not dependent on the carbon source or growth medium (**Fig S2B and Fig S2C**), and current increased after supplementation of riboflavin in the growth medium (**Fig S2D**). Collectively, these results demonstrate that *L. plantarum* possesses robust EET activity and can utilize ferric iron or an electrode as extracellular electron acceptors under a variety of conditions.

### Iron reduction in LAB is associated with the presence of *ndh2* and *pplA*

Because iron reduction by *L. monocytogenes* requires the genes in a 8.5 kb gene locus encoding a flavin-based EET (FLEET) pathway (Light et al., 2018), we examined for the presence of these genes in 1,788 complete LAB genomes deposited in NCBI. Homology searches identified the complete FLEET locus in 11 out of 38 genera including diverse LAB such as *Enterococcus* and *Lacticaseibacillus* (**Fig 2A**). The other LAB genera either lack multiple FLEET pathway genes or, as was observed for all 68 strains of *Lactococcus*, contain all genes except for *pplA*, an extracellular flavin-binding reductase. Among the lactobacilli, genomes of 19 out of 94 species contain the entire FLEET system (**Fig S3**). The lactobacilli species with the entire FLEET system are homofermentative and are distributed between different phylogenetic groups (Zheng et al., 2020). These data show that the FLEET locus is conserved across LAB genera besides *L. plantarum*, including other homofermentative LAB species known to colonize host and food environments.

**Figure 2.**
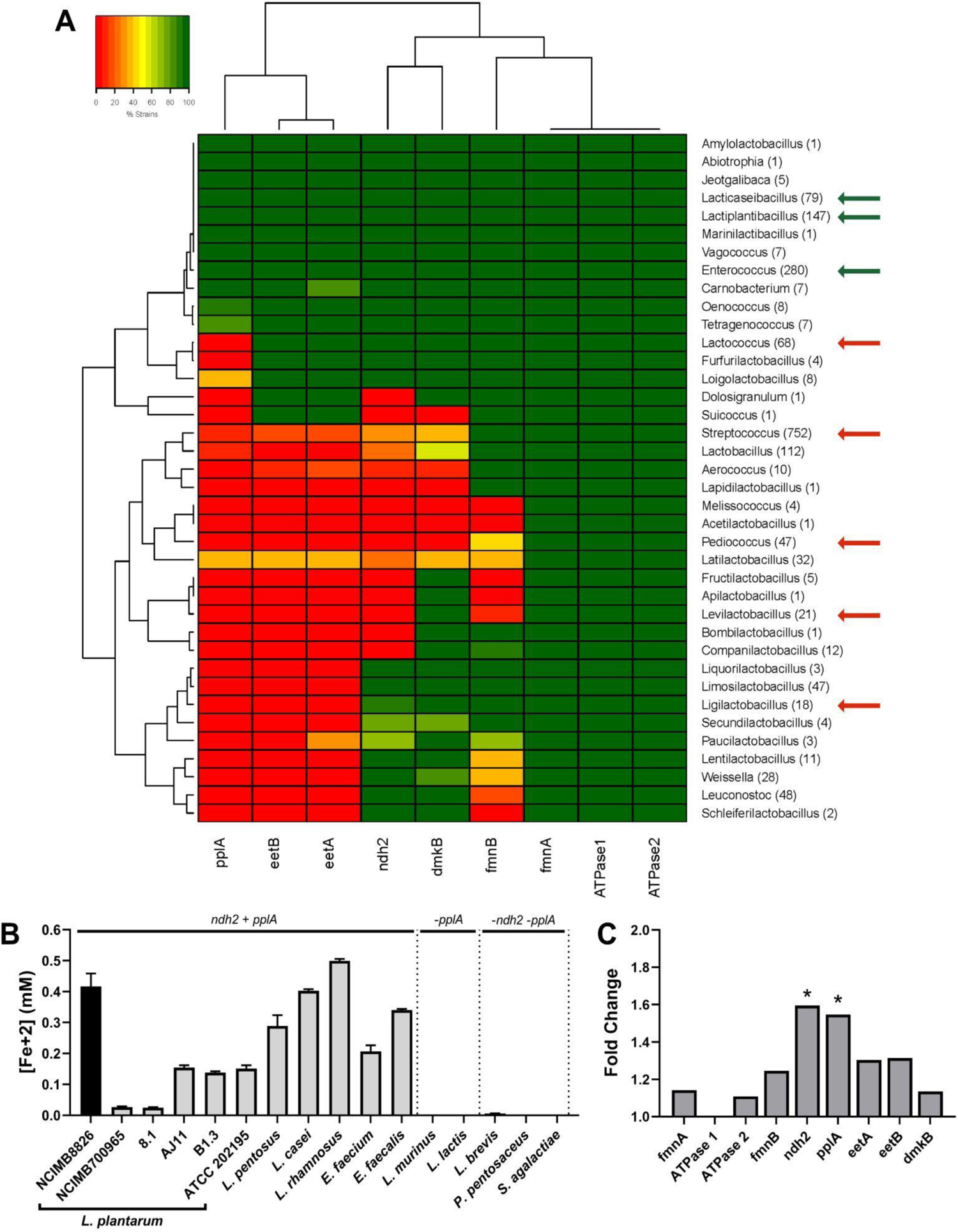
The FLEET genes ndh2 and pplA are associated with iron reduction by LAB. (A) Heatmap showing the percentage of genera in the Lactobacillales order containing FLEET genes. Homology searches were conducted using tBLASTx for 1,788 complete LAB genomes in NCBI (downloaded 02/25/2021) against the L. plantarum NCIMB8826 FLEET locus. A match was considered positive with a Bit-score > 50 and an E-value of < 10-3. Arrows designate genera tested for iron reduction activity; green = EET-active with Fe3+, red = EET-inactive with Fe3+. (B) Reduction of Fe3+ (ferrihydrite) to Fe2+ in PBS with 20 µg/mL DHNA and 55 mM mannitol. The avg ± stdev of three replicates per strain is shown. (C) Relative expression of NCIMB8826 FLEET locus genes in mMRS with 20 µg/mL DHNA and 1.25 mM ferric ammonium citrate compared to growth in mMRS. Significant differences in expression were determined by the Wald test (n = 3) with a Log2 (fold change) > 0.5 and an FDR-adjusted p-value of < 0.05.

To determine whether LAB FLEET gene presence was associated with EET activity, a diverse collection of LAB strains were examined for their capacity to reduce ferrihydrite. The assay showed that isolates of *L. plantarum*, *Lactiplantibacillus pentosus, Lacticaseibacillus rhamnosus, Lacticaseibacillus casei,* and *Enterococcus faecium* and *Enterococcus faecalis* are capable of Fe^3+^ reduction (**Fig 2B**). The genomes of those species also contain a complete FLEET pathway (**Fig 2A and Fig S3**). Conversely, strains of *Lactococcus lactis*, *Ligilactobacillus murinus*, *Levilactobacillus brevis*, *Pediococcus pentosaceus*, and *Streptococcus agalactiae* showed little to no iron reduction activity (**Fig 2B**). The presence of FLEET-associated genes varied between those species, but only those containing both *ndh2,* a membrane-bound, type-II NADH dehydrogenase, and *pplA* were able to reduce iron.

*L. plantarum* NCIMB8826 exhibited the highest EET activity resulting in at least 2.5-fold greater Fe^3+^ reduction than the other strains tested (**Fig 2B**). The *L. plantarum* NCIMB8826 genome and the genomes of 138 other strains queried also harbored a complete FLEET pathway including *ndh2* and *pplA* (**Fig S3 and Fig S4A**). Interestingly, *L. plantarum* strains NCIMB700965 and 8.1 could not reduce Fe^3+^ but possessed all genes in the FLEET system. Closer examination of both strains by aligning their FLEET loci with NCIMB8826 revealed unique IS30-family transposons in the intergenic promoter regions spanning *ndh2* and *pplA* (**Fig S4A**). These genes were minimally expressed in *L. plantarum* NCIMB700965 and 8.1 in comparison to NCIMB8826 (**Fig S4B**). *ndh2* and *pplA* were also the only two genes in the FLEET pathway that were induced when *L. plantarum* NCIMB8826 was incubated in FLEET-inducing conditions (**Fig 2C and Fig S4C**). Both *ndh2* and *pplA* were induced (∼1.6-fold, p ≤ 0.05) in MRS containing mannitol, DHNA, and ferric ammonium citrate (**Fig 3C**), but were not up-regulated when either DHNA or ferric ammonium citrate were omitted from the culture medium (**Fig S4D**). Taken together, these data show that widespread iron reduction in LAB is tightly associated with the presence and upregulation of *ndh2* and *pplA*, suggesting they are required for EET.

**Figure 3.**
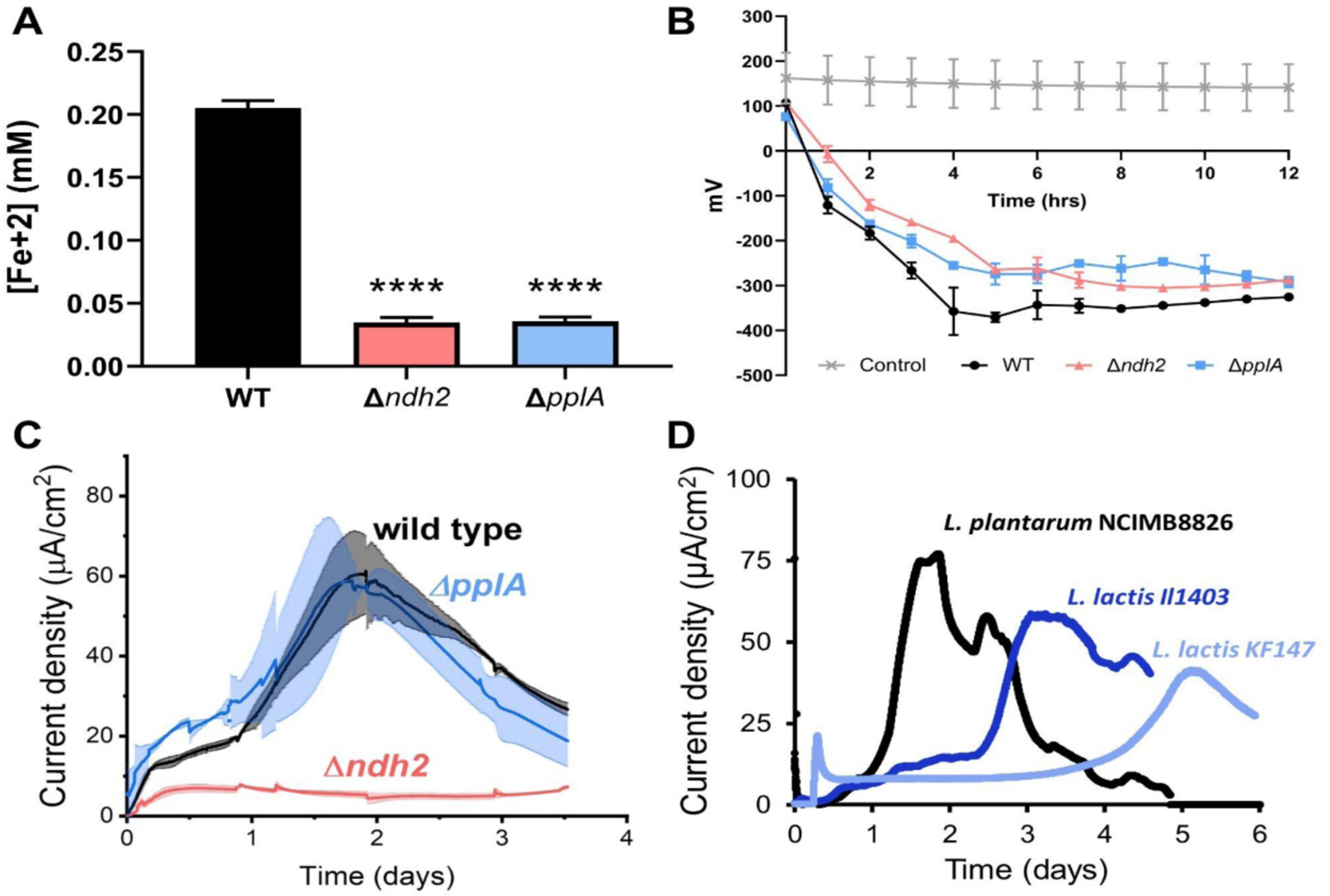
*L. plantarum* requires *ndh2* and conditionally *pplA* for EET. **(A)** Reduction of Fe^3+^ (ferrihydrite) to Fe^2+^ with wild-type *L. plantarum* or FLEET deletion mutants in the presence of 20 µg/mL DHNA and 55 mM mannitol. Significant differences determined by one-way ANOVA with Dunnett’s post-hoc test (n = 3), **** p ≤ 0.0001. **(B)** Redox potential of mMRS supplemented with 20 µg/mL DHNA and 1.25 mM ferric ammonium citrate after inoculation with wild-type *L. plantarum* or FLEET deletion mutants. **(C)** Current density generated by wild-type *L. plantarum* and deletion mutants in mannitol-CDM (mCDM) supplemented with 20 µg/mL DHNA. The avg ± stdev of three replicates is shown. **(D)** Current density generated by *L. plantarum* and two *L. lactis* strains lacking *pplA* in mCDM. *L. plantarum* was supplemented with 20 µg/mL DHNA. The data correspond to the average of two replicates per strain.

### *ndh2* is required and *pplA* is conditionally required for *L. plantarum* EET

In order to confirm the necessity of *ndh2* and *pplA* for EET in *L. plantarum*, we constructed *ndh2* and *ppA* deletion mutants in *L. plantarum* NCIMB8826. Significantly lower quantities of ferrihydrite were reduced by either mutant compared with the wild-type strain (**Fig 3A**). The loss of these genes also changed how *L. plantarum* altered extracellular redox potential (ΔmV) *in situ* in MRS culture medium. Although growth rates and cell yields were equivalent to wild-type *L. plantarum* during growth in FLEET-inducing conditions in MRS culture medium (**Fig S5A**), the rate of ΔmV reduction was lower for the Δ*ndh2* and Δ*pplA* mutants (**Fig 3B**).

Wild-type *L. plantarum* was also able to reduce more ferrihydrite than Δ*ndh2* and Δ*pplA* strains when ΔmV_max_ was reached during mid-exponential phase growth (**Fig S5B**) and this difference persisted in stationary phase cells (**Fig S5C**). These observations show that *ndh2* and *pplA* contribute to the capacity of *L. plantarum* to reduce iron and its extracellular environment.

Use of an anode as an external electron acceptor instead of ferrihydrite showed a similar, but not identical genetic dependency. *L. plantarum* Δ*ndh2* produced a significantly lower current density (**Fig 3C**) and a lower peak current (**Fig S5D**). Surprisingly, *L. plantarum* Δ*pplA* was able to produce the same amount of current as the wild-type strain, suggesting that the lipoprotein PplA is not essential and might not be involved in anode reduction through EET. This observation led us to investigate the anodic-EET ability of other LAB species lacking *pplA* like *Lactococcus lactis* (**Fig 3D**). DHNA was not provided to these strains because they can synthesize DMK and other quinones (Rezaïki et al., 2008). Both *L. lactis* strain IL1403 and strain KF147 were capable of current generation, confirming that PplA is not essential for LAB to produce current. This is consistent with the finding that other extracellular reductases besides PplA are responsible for EET activity in Gram-positive bacteria (Light et al., 2019). Taken together these results show that EET activity is dependent upon the presence of the putative FLEET locus, and specifically *ndh2* and conditionally *pplA*.

### *L. plantarum* increases energy conservation and balances intracellular redox state when performing EET

Building from studies in *E. faecalis* (Keogh et al., 2018), it has been suggested that EET improves growth by either enabling iron to be acquired as a macronutrient or by enhancing respiration (Jeuken et al., 2020). It is worth noting that several studies have shown that *L. plantarum* does not require iron to grow (Elli et al., 2000; Weinberg, 1997). To test whether EET allowed increased iron acquisition by *L. plantarum*, we measured intracellular iron by Inductively Coupled Plasma-Mass Spectrometry (ICP-MS). There was no significant difference in the amount of intracellular iron between *L. plantarum* using iron to perform EET and not performing EET (**Fig S6A**). Moreover, deletion of *ndh2* did not significantly change the amount of intracellular iron (**Fig S6B**). In contrast to studies in *E. faecalis* in which iron supplementation leads to intracellular accumulation of this metal (Keogh et al., 2018), these data show that *L. plantarum* does not use EET to increase its acquisition of iron or other metals, suggesting EET may instead play a role in energy conservation.

We next sought to understand if EET impacts energy conservation in *L. plantarum* by comparing its growth and ATP levels in the presence of a polarized anode. The highest current density (i.e., greatest EET activity) produced by *L. plantarum* in CDM typically occurred within 24 h after inoculation into the bioreactor (**Fig 1D**). At the point of maximum current, there was an approximately 4-fold higher dry cell weight and 2-fold higher numbers of viable cells compared to *L. plantarum* incubated in open circuit (OC) conditions (**Fig 4A-B**). Under the same polarized anode conditions, lag phase was shortened (**Fig 4C**), indicating that EET allowed *L. plantarum* to start growth earlier. Besides an earlier initiation of growth, intracellular ATP levels at peak current production were significantly higher (4.5-fold) under EET compared to OC conditions (**Fig 4D**) (**Table 1**). ATP levels were also greater in *L. plantarum* when in the presence of both mannitol and DHNA, compared to either mannitol or DHNA separately (**Fig 4D**). Thus, EET allows *L. plantarum* to initiate growth more rapidly and accumulate more ATP, indicating that EET significantly increases energy conservation in *L. plantarum*.

**Figure 4.**
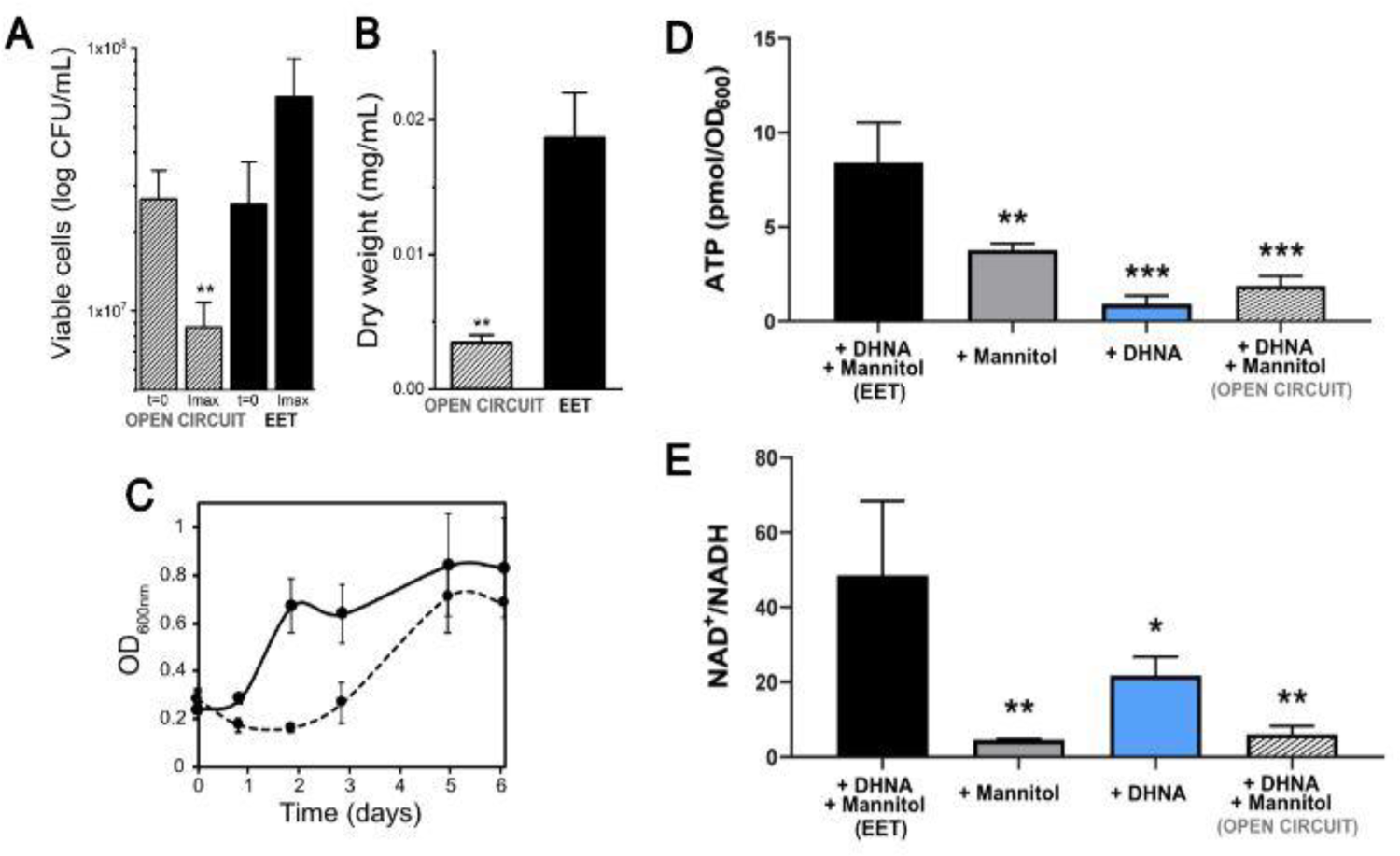
Growth, ATP, and redox balance of *L. plantarum* changes when an anode is provided as an extracellular electron acceptor. **(A)** Viable cells and **(B)** dry weight at the point of maximum current density from **Fig 2B** under current circulating conditions (EET) and at open circuit conditions (OC) at the same time point. **(B)** Change in cell numbers measured by OD_600nm_ overtime in the bioreactors under EET and OC conditions. **(D)** ATP production per OD unit and **(E)** NAD^+^/NADH ratios at the point of maximum current density. The bioreactors were shaken vigorously to dislodge cells before sampling. The avg ± stdev of three replicates is shown. Significant differences were determined by one-way ANOVA with **(A and B)** Dunn-Sidak post-hoc test (n = 3) and **(D and E)** Dunnett’s post-hoc test (n = 3), * p < 0.05; ** p < 0.01; *** p < 0.001; **** p < 0.0001.

**Table 1.** Bioenergetic balances suggest energy conservation under EET conditions occurs via substrate level phosphorylation. The reactors contained 20 ug/mL of DHNA and mannitol as electron donor.

|  | NADH generated* | NADH consumed* | Calculated ATP** (from metabolites) | Biomass yield | Y<sub>fermentation</sub> | Y<sub>mannitol</sub> | Y<sub>ATP</sub> |
| --- | --- | --- | --- | --- | --- | --- | --- |
| Units | mM | mM | mM | g-dw/mol-mannitol | (mmol product/mmol-mannitol) | mol ATP/mol mannitol | g dw/mol ATP |
| **EET** | 31.5 | 6.44±0.48 via anode<br>15.10±2.6 via SLP | 15.87±1.71 | 4.85±0.33 | 1.26±0.12 | 1.51±0.16 | 3.24±0.43 |
| **OC** | 19.1 | 5.51±0.97 via SLP | 4.7±0.6 | 7.21±1.41 | 0.72±0.09 | 0.74±0.09 | 9.64±1.04 |

\*Calculated on the basis of production of 3 mol of NADH produced per mol of mannitol and, 1 mol of NAD<sup>+</sup> per lactate, 2 mol of NAD<sup>+</sup> per ethanol and 2 mol of NAD<sup>+</sup> mol per succinate produced and 0.5 mmol of NAD<sup>+</sup> per mol of electrons harvested on the anode
\*\* Calculated on the basis of production of 1 mol of ATP per lactate, 2 mol per acetate, 1 mol per ethanol and 3 mol per succinate produced.

Because fermentation, anaerobic respiration, and aerobic respiration are each associated with a different NAD^+^/NADH ratio, energy conservation is linked to intracellular redox homeostasis (Holm et al., 2010). Therefore, we probed redox homeostasis in *L. plantarum* under EET conditions by measuring intracellular NAD^+^/NADH at the point of maximum current density (**Fig 4E**). *L. plantarum* showed a 8-fold higher NAD^+^/NADH ratio under EET conditions compared to OC (**Fig 4E**). This result was not limited to the presence of a polarized anode as *L. plantarum* also contained significantly a higher NAD^+^/NADH ratio when ferrihydrite was available as a terminal electron acceptor (**Fig S7**). These NAD^+^/NADH ratios are more similar to those found for in *E. coli* performing aerobic respiration (de Graef et al., 1999) or *G. sulfurreducens* performing anaerobic respiration than in LAB performing fermentation (Guo et al. 2017). Taken together, our data strongly suggest that EET is involved in energy conservation, and the intracellular redox balance during EET mimics a respiratory rather than a fermentative process.

### EET increases fermentative metabolism through substrate level phosphorylation and reduction in extracellular pH

Metal-reducing bacteria use EET in anaerobic respiration (Richter et al., 2012; Shi et al., 2007). *L. plantarum* can perform anaerobic respiration in the presence of exogenous menaquinone and heme (Brooijmans et al., 2009), suggesting that EET may be used during anaerobic respiration to increase energy conservation. To test this hypothesis, we examined whether electron transfer proteins needed for PMF generation aerobic and anaerobic respiration are required for *L. plantarum* EET. Neither the addition of heme to restore bd-type cytochrome (*cydABCD*) used in aerobic respiration, nor deletion of the respiratory nitrate reductase (Δ*narGHJI*) significantly altered current production (**Fig S9A-B**). Because Ndh2 is a type-II NADH dehydrogenase which does not contribute to a proton gradient (Lin et al., 2008; Nakatani et al., 2020), these observations show that EET does not involve the known PMF-creating proteins in *L. plantarum*. Since respiration is associated with the tricarboxylic acid (TCA) cycle, we also examined production of succinate which is the terminal end-product of the reductive branch of this pathway in *L. plantarum* (Tsuji et al., 2013). EET did not increase the succinate concentration (**Fig S8**), indicating that EET did not cause additional metabolic flux through its TCA cycle. Thus, our results suggest that anaerobic respiration is not responsible for increased energy conservation under EET conditions.

An alternative hypothesis is that increased energy conservation under EET conditions is driven by increases in fermentation. Known as a primarily fermentative organism, *L. plantarum* uses glycolysis to convert mannitol to two molecules of pyruvate which is then converted to lactate or ethanol via NADH-consuming steps, or acetate via an ATP-generating reaction using substrate-level phosphorylation (Dirar and Collins, 1972). With a polarized anode, the yield of fermentation products (acetate, lactate and ethanol) per cell was 2.6-fold higher compared to OC conditions (**Fig 5A**). The culture medium pH was also significantly lower than under OC (**Fig 5B**). A similar acidification of the medium was observed for Δ*pplA*, but not for Δ*ndh2*, when an anode was present as electron acceptor, indicating that current production capacity is key to drop the pH (**Fig S5E**). These results show that EET allows *L. plantarum* to ferment and to acidify the medium to a greater degree.

**Figure 5.**
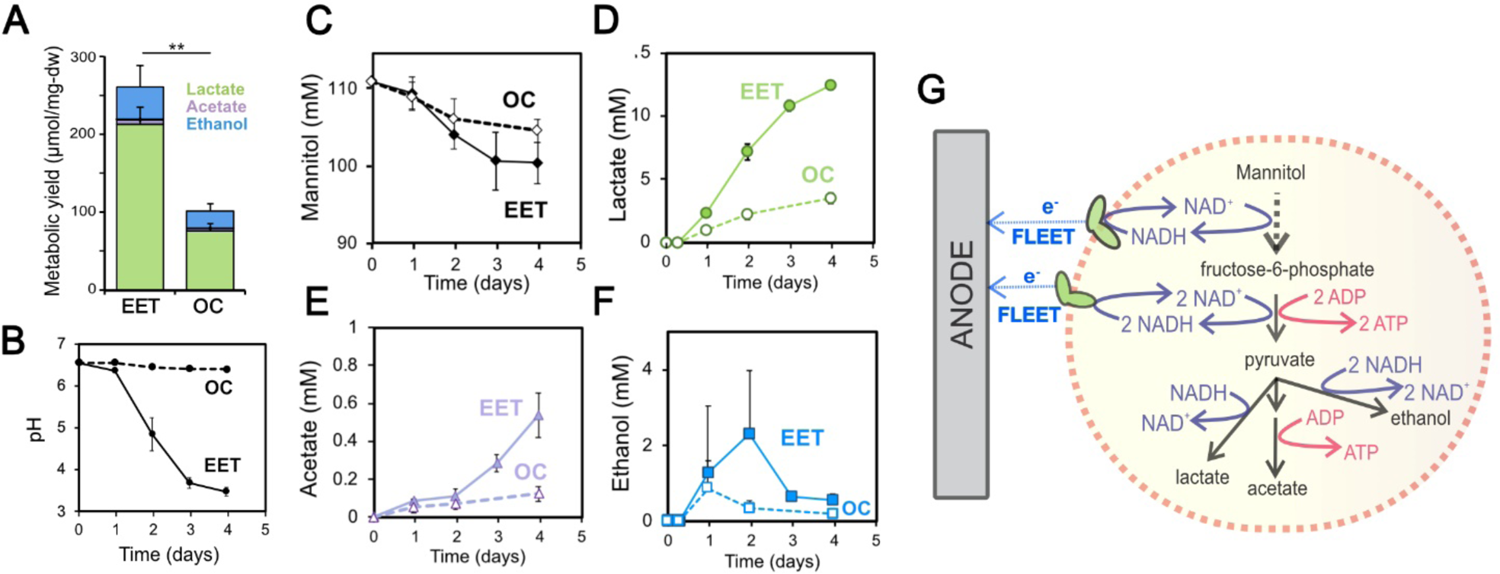
Fermentation fluxes are increased when an anode is provided as an extracellular electron acceptor. **(A)** Metabolic yields of *L. plantarum* end-fermentation products under open circuit conditions (OC) and current circulating conditions (EET) in mCDM supplemented with 20 µg/mL DHNA. **(B)** pH measurements and **(C)** mannitol, **(D)** lactate, **(E)** acetate and **(F)** ethanol concentrations over time under OC and EET conditions. **(G)** Schematic of proposed model for NADH regeneration during fermentation of mannitol in the presence of an anode as electron sink for *L. plantarum*. The avg ± stdev of three replicates is shown. Significant differences were determined by One-way ANOVA with Dunn-Sidak post-hoc (n = 3), ** p ≤ 0.01.

We also observed that EET led to higher cellular metabolic fluxes (change of substrate over time). *L. plantarum* utilized more mannitol and produced more acetate and lactate under EET conditions (**Fig 5C-F**). Cells performing EET were overall 2-fold faster at consuming mannitol (**Fig 5C**) compared to OC conditions by day four. Mannitol consumption increased after one day, approximately when the cells exited lag phase (**Fig 4C**), suggesting that growth drove that consumption. Although the final OD_600nm_ and dry cell weight were not significantly different (**Table S1**), the overall rates of acetate and lactate production also increased 3.4 and 3.6 times (**Fig 5D-E**), respectively. EET also allowed cells to produce more fermentation products per each mol of mannitol utilized (Y_fermentation_, **Table 1**). These data indicate that EET increases the flux and final yield of fermentation in *L. plantarum*.

Because the production of acetate yields ATP, these results also suggested that the increase in ATP generation under EET conditions may be due to substrate-level phosphorylation. To probe whether EET-associated increase in fermentative flux could account for the changes in ATP generation, we calculated fermentation balances (**Table 1**). The concentrations of fermentation products (**Table S1**) were used to estimate the total ATP in the presence (ATP_EET_) and absence of EET (ATP_OC_). The estimated ATP_EET_ was 3-fold higher than ATP_OC_ (**Table 1**), a result that is consistent with the higher accumulation of ATP observed (**Fig 4D**). This analysis shows quantitatively that the increase in ATP generation with EET is due to an increase in fermentative flux and substrate-level phosphorylation.

### EET shifts how *L. plantarum* uses electron acceptors and converts ATP into biomass

Thus far, our results provided an unusual picture of the energy metabolism of *L. plantarum* under EET conditions; while EET significantly shifted the intracellular redox state to a more respiratory-like balance, its increased ATP yield was mainly accounted for by an increased fermentative flux and yield. Another major difference in fermentation and anaerobic respiration is the use of the intracellular organic versus inorganic electron acceptors. To more deeply understand how *L. plantarum* uses organic molecules and the anode as electron acceptors when performing EET, the electron balances under EET and OC conditions were calculated (**Table 1** and **Table S1**). We estimated the NADH produced from the mannitol consumed and both the NADH reoxidized through the reduction of the anode (measured as current) and through the production of metabolites via NADH-consuming pathways (measured as lactate, ethanol and succinate). This allowed us to obtain a global balance in the NAD^+^/NADH ratio.

Under OC conditions, the concentrations of lactate, ethanol and succinate produced indicated that only about a third of the NADH produced from the oxidation of mannitol to pyruvate was reoxidized to NAD^+^ (5.1 mM NADH reoxidized vs. 19.1 mM NADH generated, **Table 1**), qualitatively agreeing with the low NAD^+^/NADH ratios measured (**Fig 4E**). In contrast, electron balance calculations showed that more than two-thirds of the NADH was reoxidized under EET conditions (21.5 mM NADH consumed vs. 31.5 mM generated, **Table 1**), a result that is consistent with the significantly higher NAD^+^/NADH ratios measured (**Fig 4E**). Interestingly, it was estimated that 48% of the total NADH generated was oxidized through fermentation, while 20% of the NADH was oxidized using the electrode as a terminal electron acceptor (**Table 1**). Thus, *L. plantarum* growing under EET conditions achieves a more oxidized intracellular redox balance by both more completely fermenting mannitol to lactate and ethanol and by using the electrode as a terminal electron acceptor (**Fig 5G**). These observations reinforce that the energy metabolism of *L. plantarum* under EET conditions utilizes elements of both fermentation and anaerobic respiration.

To connect this energy conservation with growth, the ATP requirements to grow biomass for each condition (Y_ATP_) were estimated using the calculated ATP and the measured dry weight. Under OC conditions, the Y_ATP_ obtained (9.64 ± 1.04 g dw/mol ATP) for *L. plantarum* was similar to that observed previously (10.9 g-dw/mol ATP) (Dirar and Collins, 1972). Hence, without EET, the ATP generated from fermentation was converted into biomass at the expected efficiency. In contrast, a significantly lower Y_ATP_ was reached for *L. plantarum* performing EET (3.24 ± 0.43 g dw/mol ATP) (**Table 1**). This observation indicates that under EET conditions, either more ATP is required to accumulate biomass or ATP-accumulating mechanisms are stimulated (also known as energy spilling) (Russell, 2007). EET conditions also resulted in 74% more ATP per mol of fermented mannitol (Y_mannitol_). Consequently, molar biomass yields (g-dw/mol-mannitol) under EET conditions were significantly lower (**Table 1**), in agreement with previous observations in respiratory electroactive species (Esteve-Núñez et al., 2005). These calculations show that when *L. plantarum* performs EET, it more efficiently produces ATP, but this ATP is less efficiently utilized to make biomass. Overall, these results show an intriguing pattern of ATP generation and its use to form biomass that further suggests the existence of a different energy conservation strategy when the EET chain is activated.

### EET is active in vegetable fermentations

Our previous results inspired us to explore whether EET could occur in a physiological niche of LAB such as fermented foods. LAB are necessary for the making of many fermented fruit and vegetable foods and the properties of those foods depend on the metabolic diversity of the LAB strains present (Marco and Golomb, 2016). Plant tissues also contain a much wider variety of carbon substrates and potential electron acceptors than the CDM used in our prior experiments. To study the physiological and biotechnological relevance of EET in food fermentations, kale juice was fermented using *L. plantarum* as a starter culture (**Fig 6A**). The fermentation of kale juice was measured under EET conditions (a polarized anode with or without DHNA), and an OC control (a nonpolarized anode with DHNA) to establish the role of DHNA and electrical current circulation on the fermentation process. An additional bioreactor without cells, but with DHNA, was operated to identify any possible electrochemical-driven conversion of substrates. When *L. plantarum* was added to the prepared kale juice, approximately 10-fold more current was generated during EET conditions with DHNA (EET+DHNA) as compared to abiotic and biological non-EET promoting conditions (no DHNA) (**Fig 6B**). This current was comparable to the current generated in laboratory medium (**Fig 1D**), indicating that robust EET by LAB is possible in the complex physiological conditions of a food fermentation.

**Figure 6.**
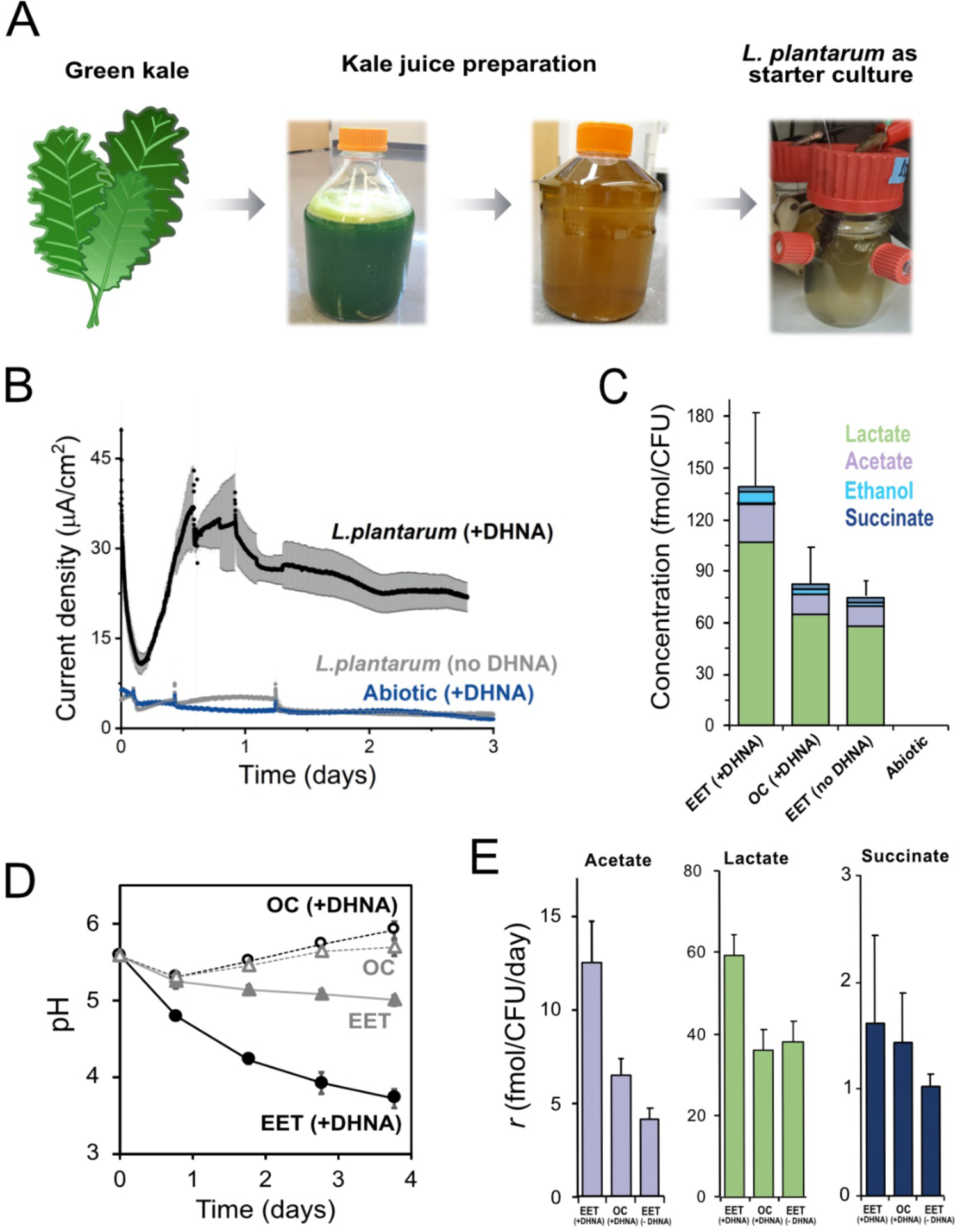
EET in a kale juice increases the production of fermentation end products. **(A)** Preparation of kale juice medium used for fermentation in bioelectrochemical reactors. **(B)** Current density production measured from kale juice medium over time in the presence of *L. plantarum* and 20 µg/mL DHNA, no DHNA, or under abiotic conditions with addition of 20 µg/mL DHNA. The anode polarization was maintained at 0.2 V_Ag/AgCl_. **(C)** Normalized total quantities of the metabolites detected per cell (CFU_max_ used for calculations). **(D)** pH measurements over time under different conditions tested on a second set of kale juice fermentations performed under the same conditions. (**E)** Production rate per viable cell, *r*, of lactate, acetate, and succinate. The avg ± stdev of three replicates is shown.

We next investigated the impact of EET on *L. plantarum* growth and metabolism in the kale juice fermentation. Significant changes in the pH and fermentation products were detected under EET conditions (**Fig 6C-D**). These differences occurred in the absence of significant changes in viable cell numbers (**Fig S10A**) at the time points measured. As previously observed using laboratory culture media, an approximately two-fold greater accumulation of total end-fermentation products per cell was obtained when cells interacted with an anode in the presence of DHNA (**Fig 6C**). In the kale juice fermentation, EET+DHNA conditions enhanced both lactate and acetate production per cell (**Fig 6E and S10B**). Thus, when DHNA was provided, EET enhanced the overall yield of fermentation end-products and their production rates per cell, mimicking our observations in laboratory media (**Fig 5C-D**). EET also led to a significantly higher acidification of the kale juice compared to OC conditions, and the presence of DHNA dramatically enhanced this pH drop (**Fig 6D**). In general, when no DHNA was supplied but an anode was present as an electron acceptor, the fermentation process was very similar to OC conditions. This means in kale juice a source of quinones is essential to support *L. plantarum* EET activity. Overall, these results show that EET under physiological conditions impacts cellular metabolism in *L. plantarum* by increasing metabolic flux which ultimately can affect the flavor profile of fermented foods (Chen et al., 2017).

## DISCUSSION

Increases in fermentation and energy conservation from EET have important bioenergetic implications for the mainly fermentative LAB. We showed that *L. plantarum* and other diverse LAB species perform EET if flavins and quinones are present. EET activity by *L. plantarum* is robust, as it results in current densities comparable to those reported for respiratory metal-reducing bacteria. EET allows *L. plantarum* to balance the intracellular redox and generate a high NAD+/NADH ratio. This metabolic impact increases fermentation yield and flux, shortens lag phase, and increases ATP production. *L. plantarum* EET activity requires an NADH dehydrogenase (Ndh2) and conditionally requires an extracellular, flavin-binding reductase (PplA). Thus, EET in *L. plantarum* is a hybrid energy metabolism that contains metabolic features of fermentation and redox features of anaerobic respiration. This pathway is active in *L. plantarum* in physiologically relevant conditions and results in an increased metabolic flux and acidification rate.

### The combined EET fermentation hybrid metabolism is distinct from anaerobic respiration, fermentation, and other energy conservation strategies

When performing EET, the metabolism of *L. plantarum,* a primarily fermentative bacterial species, is fundamentally different from EET-driven, anaerobic respiration of metal-reducing bacteria or anaerobic respiration of LAB. Although aspects of EET in *L. plantarum* and metal-reducing *Geobacter* spp. are similar, such as the upregulation of NADH dehydrogenase, the reduction rate of extracellular electron acceptors, and the high NAD^+^/NADH ratio, other metabolic features stimulated by EET in these two organisms are starkly different (see comparison in **Table S7**). *L. plantarum* regenerates a substantial fraction of its NADH by directing metabolic flux through fermentative pathways to achieve a higher ATP yield. By comparison, *Geobacter* spp. direct its metabolic flux through the TCA cycle, relies almost exclusively on extracellular electron acceptors to regenerate NADH, and produces ATP exclusively through oxidative phosphorylation. Comparing EET and respiratory metabolism in LAB (**Table S6**) shows that both metabolisms require quinones, but respiration also requires heme. Our findings and similar findings in *E. faecalis* (Pankratova et al., 2018) confirm that heme is not required for EET. Moreover, EET also differs from respiration in LAB because it occurs at the start of or prior to exponential phase growth, does not change the final cell density, and increases fermentation with no effect on the resultant proportions of lactate, acetate, and ethanol (Duwat et al., 2001). Thus, EET in LAB diverges from respiration in metal-reducing bacteria or LAB in its metabolic pattern and energetic consequences.

This EET mechanism is also a novel energy generation strategy compared to known forms of fermentative metabolism in LAB (comparison in **Table S6**). *L. plantarum* and other LAB reduce alternative intracellular electron acceptors like citrate, fructose, and phenolic acids, resulting in increased intracellular NAD^+^/NADH ratios (Hansen, 2018). Unlike these examples, however, the reduction of extracellular Fe^3+^ or an anode by EET requires an electron transport chain and the shuttling electrons outside of the cell. In addition, the reduction of the oxygen and organic compounds for cofactor regeneration by LAB leads to a metabolic shift towards acetate production and altered metabolic end-product ratios (Gänzle, 2015). These differences show how the combined EET fermentation hybrid metabolism is distinct from other pathways that alleviate reduced intracellular conditions in LAB.

Previous studies have reported a simultaneous use of fermentation and electron transport elements, such as in the so-called, respiro-fermentation in *Saccharomyces cerevisiae* (Blom et al., 2000-5). However, respiro-fermentation produces ATP and maintains intracellular redox balance through substrate-level phosphorylation and/or oxidative phosphorylation using separate pathways. Our data strongly suggests a single pathway is responsible for both ATP generation and intracellular redox balance. This hybrid fermentation mode is also different from the electron bifurcating mechanism, in which the extra ATP generation is driven by the creation of a H^+^ or Na^+^ potential from the oxidation of a ferredoxin (Buckel and Thauer, 2013). Another poorly understood example of the use of substrate-level phosphorylation and electron transport chains to balance intracellular redox states is found in the non-fermentative bacterium *S. oneidensis*.

Although this species relies on SLP to grow anaerobically, it couples the reduction of fumarate, an electron acceptor, to ATP synthesis via the creation of PMF (Hunt et al., 2010). Unlike in these examples, EET in LAB does not involve PMF creating elements and the FLEET chain drives ATP generation through substrate-level phosphorylation. Overall, *L. plantarum* EET-associated metabolism contains features of both fermentation (e.g., substrate-level phosphorylation, substrates utilized) and respiratory metabolisms (e.g., NAD^+^/NADH ratios, NADH dehydrogenase required) (**Table S6**). Thus, to our knowledge, the strategy that *L. plantarum* uses to generate ATP performing EET constitutes a novel mode of energy conservation.

### Different mechanisms of EET appear to be widespread in LAB

Based on our observations and others, we propose that EET is widespread in LAB and occurs by different mechanisms. Besides *L. plantarum*, we showed that *L. lactis* is able to generate current, despite lacking *pplA*. Current generation by *L. lactis* was observed previously, found to be riboflavin dependent, and resulted in a small metabolic, yet to be defined, shift in which the flux through NADH-oxidizing pathways was reduced and ATP generating pathways were increased (Freguia et al., 2009; Masuda et al., 2010). Extracellular reduction of tetrazolium violet is further evidence of EET activity in *L. lactis* and this activity was dependent on the presence of both quinones and an NADH dehydrogenase (NoxAB) (Tachon et al., 2009). *E. faecalis* is another LAB that performs EET and similar to *L. plantarum* requires quinones (Pankratova et al., 2018) and a type-II NADH dehydrogenase (Ndh3) (Hederstedt et al., 2020) for Fe^3+^ reduction. In contrast to this mechanistic similarity, *E. faecalis* performs EET using matrix-associated iron resulting in both increased final cell biomass and intracellular iron (Keogh et al. 2018). Moreover, unlike *L. plantarum* and *L. lactis*, the presence of PplA is necessary for anode reduction, but not Fe^3+^ reduction (Hederstedt et al., 2020). The conditional need for PplA in EET may be explained by the identification of other flavin-binding, extracellular reductases amongst Gram-positive organisms, such as FrdA (acting on fumarate) in *L. monocytogenes* and UrdA (acting on urocanate) in *Enterococcus rivorum* (Light et al., 2019). Thus, there may exist a yet unidentified extracellular reductase in *L. plantarum* and *L. lactis* required for anode reduction. Thus, our findings substantially expand the understanding of this capacity by elucidating a new pattern of metabolic changes associated with EET. It seems likely that these many mechanisms reflect the ability of EET to alleviate constraints of intracellular redox balance in fermentative metabolism across LAB.

### EET has important implications for ecology and biotechnology of LAB

Conservation of the FLEET pathway among different LAB species supports the premise that this hybrid fermentation with EET provides an important metabolic strategy for these bacteria in their natural habitats. LAB with a complete FLEET system are homofermentative, meaning that lactate is the sole or major product of metabolism, thus underscoring the distinct ways homofermentative and heterofermentative LAB have evolved for energy conservation (Salvetti et al., 2013). *L. plantarum* and other LAB with FLEET systems such as *L. casei* are genetically and metabolically diverse and grow in a variety of nutrient rich environments including dairy and plant foods and the digestive tract (Cai et al., 2009; Martino et al., 2016; Siezen et al., 2010). Those environments also are rich sources of sources of quinones, flavins, and extracellular electron acceptors (such as iron) (Cataldi et al., 2003; Fenn et al., 2017; Kim, 2017-9; Roughead and McCormick, 1990; Walther et al., 2013). Increased organic acid production and environmental acidification by LAB with this hybrid metabolism would provide an effective mechanism to inhibit competing microorganisms and confer a competitive advantage for growth. The increased ATP relative to biomass generation observed during growth on mannitol might also give sufficient readiness for using this energy later on to outcompete neighboring organisms (Russell and Cook, 1995). These effects of EET may be particularly important on plant tissues and intestinal environments, wherein LAB tend to be present in low numbers. Besides our observation that *L. plantarum* performs EET in kale juice, the FLEET pathway is important for intestinal colonization by both *L. monocytogenes* (Light et al., 2018) and *E. faecalis* (Lam et al., 2019), and the *L. plantarum* FLEET genes were highly induced in the small intestine of rhesus macaques (Golomb et al., 2016).

The hybrid fermentation metabolism of LAB also has technological relevance. For many LAB food fermentations, acidification of the food matrix is required to prevent the growth of undesired microorganisms and result in a more consistent and reproducible product. Starter cultures are frequently selected based on their capacity for rapid growth and acid production (Bintsis, 2018). In the presence of the anode, exposure of *L. plantarum* to EET conditions during kale juice fermentation increased the acidification rate. Thus, this shows that EET metabolism is active in complex nutritive environments such as kale leaf tissues that contain other potential electron acceptors besides the anode and diverse carbon sources (glucose, fructose, sucrose) (Thavarajah et al., 2016). This example also shows how electro-fermentation, the technological process by which fermentation is modulated using electrodes, can be used to control food fermentations (Moscoviz et al., 2016; Schievano et al., 11 2016). Because *L. plantarum* also increased fermentation flux when the electrode was available as an electron sink, higher quantities of organic acid flavor compounds were formed. Therefore, by the manipulation of extracellular redox potential, electro-food fermentations may be used to control microbial growth. This would allow the creation of new or altered sensory profiles in fermented foods, such as through altered organic acid production and metabolism or synthesis of other compounds that alter food flavors, aromas, and textures (Bintsis, 2018).

### Final Perspective

We expect that the contributions of our study will improve the current understanding of energy conservation in electroactive LAB fermentations and contribute to establishing its ecological relevance. This will ultimately allow the use of EET to electronically modulate the flavor and textural profiles of fermented foods and also to expand its use to biomedicine and bioenergy (Moscoviz et al., 2016). The identification of the precise components and bioenergetic basis involved in *L. plantarum* EET will be key to unravel physiological and ecological questions and to develop other biotechnological applications.

## MATERIALS AND METHODS

### Strains and culture conditions

All strains and plasmids used in this study are listed in **Table S1**. Standard laboratory culture medium was used for routine growth of bacteria as follows: *Lactiplantibacillus* spp., *Lacticaseibacillus* spp., *Levilactobacillus brevis*, *Ligilactobacillus murinus*, and *Pediococcus pentosaceus*, MRS (BD, Franklin Lakes, NJ, USA); *Lactococcus lactis* and *Streptococcus agalactiae*, M17 (BD) with 2% w/v glucose; *Enterococcus faecalis*, and *Enterococcus faecium*, BHI (BD); and *Escherichia coli*, LB (Teknova, Hollister, CA, USA). Bacterial strains were incubated without aeration except for *E. coli* (250 RPM) and at either 30 or 37 °C. Where indicated, strains were grown in filter-sterilized MRS (De MAN et al., 1960) lacking beef extract with either 110 mM glucose [gMRS] or 110 mM mannitol [mMRS], or a chemically defined minimal medium (**Table S2**) with 125 mM glucose [gCDM] or 125 mM mannitol [mCDM] for 18 h (Aumiller et al., 2021)=. Culture medium was supplemented with 20 µg/mL of the quinone 1,4-dihydroxy-2-naphthoic acid (DHNA) (Alfa Aesar, Haverhill, MA, USA), 1.25 mM ferric ammonium citrate (VWR, Radnor, PA, USA), or 5 µg/mL erythromycin (VWR).

### DNA sequence analysis

The FLEET gene locus of *L. plantarum* NCIMB8826 was identified using NCBI basic local alignment search tool (BLAST) (McGinnis and Madden, 2004) using the *L. monocytogenes* 10403S FLEET genes (lmo2634 to lmo2641) as a reference. *L. plantarum* genes were annotated based on predicted functions within the FLEET system (Light et al., 2018). FLEET locus genes were identified in other LAB by examining 1,788 complete Lactobacillales genomes available at NCBI (downloaded 02/25/2021). A local BLAST (ver 2.10.1) database containing these genomes was queried using tBLASTx with NCIMB8826 FLEET genes a reference. A gene was considered to be present in the Lactobacillales strain genome if the Bit-score was ≥ 50 and the E-value was ≤ 10^-3^ (Pearson, 2013). Heatmaps showing the percentage of strains in Lactobacillales genera and the *Lactobacillus*-genus complex (Zheng et al., 2020) identified to contain individual FLEET genes were visualized using the R-studio package ggplot2 (Wickham, 2011) with clustering done through UPGMA. The FLEET loci of *L. plantarum* strain 8.1 and NCIMB700965 were aligned to the NCIMB8826 genome in MegAlign Pro (DNAstar Inc., Madison, WI, USA).

### Ferrihydrite reduction assays

*L. plantarum* and other strains were incubated in mMRS containing DHNA (20 µg/mL) and ferric ammonium citrate (C_6_H_8_FeNO_7_) (1.25 mM) for 18 h. Cells were collected by centrifugation at 10,000 g for 3 min, washed twice in phosphate-buffered saline (PBS), pH 7.2 (http://cshprotocols.cshlp.org), and adjusted to an optical density (OD) at 600 nm (OD_600nm_) of 2 in the presence of 2.2 mM ferrihydrite (Schwertmann and Fischer, 1973; Stookey, 1970) and 2 mM ferrozine (Sigma-Aldrich). Where indicated, 55 mM glucose or mannitol and 20 µg/mL DHNA were added. After 3 h incubation at 30 °C, the cells were collected by centrifugation at 10,000 g for 5 min and the absorbance of the supernatant was measured at 562 nm with a Synergy 2 spectrophotometer (BioTek, Winooski, VT, USA). Quantities of ferrihydrite reduced were determined using a standard curve containing a 2-fold range of FeSO_4_ (Sigma-Aldrich) (0.25 mM to 0.016 mM) and 2 mM ferrozine. The FeSO_4_ was dissolved in 10 mM cysteine-HCl (RPI, Mount Prospect, IL, USA) to prevent environmental re-oxidation of Fe^2+^ to Fe^3+^ in the standard curve.

### *L. plantarum* mutant construction

*L. plantarum* NCIMB8826 *ndh2*, *pplA*, and *narG* deletion mutants were constructed by double-crossover homologous recombination. For mutant construction, upstream and downstream flanking regions of these genes were amplified using the A/B and C/D primers, respectively, listed in **Table S3**. Splicing-by-overlap extension (SOEing) PCR was used to combine PCR products as previously described (Heckman and Pease, 2007). PCR products were digested with restriction enzymes EcoRI, SacI, SacII, or SalI (New England Biolabs, Ipswich, MA, USA) for plasmid ligation and transformation into *E. coli* DH5α. The resulting plasmids were then introduced to *L. plantarum* NCIMB8826 by electroporation. Erythromycin-resistant mutants were selected and confirmed for plasmid integration by PCR (see **Table S3** for primers). Subsequently, deletion mutants were identified by a loss of resistance to erythromycin, PCR (see **Table S3** for primers) confirmation, and DNA sequencing (http://dnaseq.ucdavis.edu).

### Bioelectrochemical reactors (BES) construction, operation, and electrochemical techniques

*L. plantarum* NCIMB8826 strains were grown overnight (approx. 16-18 h) from glycerol stocks in MRS. Cells were harvested by centrifugation (5200 g, 12 min, 4 °C) and washed twice in PBS. When *L. plantarum* wild-type EET activity versus the Δ*ndh2* (MLES100) and Δ*pplA* (MLES101) deletion mutants was compared, cells were grown as described and the number of cells was normalized across the three strains prior to inoculation in the BES. The bioreactors consisted of double-chamber electrochemical cells (Adams & Chittenden, Berkeley, CA) (**Figure 1B**) with a cation exchange membrane (CMI-7000, Membranes International, Ringwood, NJ) that separated them. A 3-electrode configuration was used consisting of an Ag/AgCl sat KCl reference electrode (BASI, IN, USA), a titanium wire counter electrode, and a 6.35-mm-thick graphite felt working electrode (anode) of 4×4 cm (Alfa Aesar, MA, USA) with a piece of Ti wire threaded from bottom to top as a current collector and connection to the potentiostat. We used a Bio-Logic Science Instruments (TN, USA) potentiostat model VSP-300 for performing the electrochemical measurements (chronoamperometry). The bioreactors were sterilized by filling them with ddH_2_O and autoclaving at 121 °C for 30 min. The water was then removed and replaced with 150 mL of filter sterilized mMRS or mCDM media for the working electrode chamber, and 150 mL of M9 medium (6.78 g/L Na₂HPO₄, 3 g/L KH_2_PO_4_, 0.5 g/L NaCl, 1 g/L NH_4_Cl) (BD) for the counter electrode chamber. Both media were supplemented with 20 µg/mL DHNA diluted 1:1 in DMSO:ddH_2_O where appropriate. To test the role of *bd*-cytochrome, heme was added in a final concentration of 10 µg/mL (diluted 1:1 in DMSO: ddH_2_O). The medium in the working electrode chamber was continuously mixed with a magnetic stir bar and N_2_ gas was purged to maintain anaerobic conditions for the course of the experiment. The applied potential to the working electrode was of +0.2 V versus Ag/AgCl (sat. KCl) (BASI, IN, USA). Reactors run under OC conditions were similarly assembled but kept at open circuit and used as control for non-current circulating conditions. Once the current stabilized, the electrochemical cells were inoculated to a final OD_600_ of 0.12-0.15 with the cell suspensions prepared in PBS. Current densities are reported as a function of the geometric surface area of the electrode (16 cm^2^). The bioreactors were sampled by taking samples under sterilized conditions at different time points for subsequent analysis. The samples for organic acids analyses were centrifuged (15,228 g, 7 min) and the supernatant was separated for High-Performance Liquid Chromatography (HPLC) assessments. Samples for ATP and NAD^+^/NADH analyses were flash frozen in a dry ice/ethanol bath.

### Metabolite analysis

Organic acids, ethanol and sugar concentrations were measured by HPLC (Agilent, 1260 Infinity), using a standard analytical system (Shimadzu, Kyoto, Japan) equipped with an Aminex Organic Acid Analysis column (Bio-Rad, HPX-87H 300 x 7.8 mm) heated at 60 °C. The eluent was 5 mM of sulfuric acid, used at a flow rate of 0.6 mL min^-1^. We used a refractive index detector 1260 Infinity II RID. A five-point calibration curve based on peak area was generated and used to calculate concentrations in the unknown samples.

### BES biomass growth determination

Bioreactors were shaken to remove the cells attached to the working electrode and afterwards sampled to measure viable cells (colony forming units (CFUs)) and total biomass (dry weight). Samples for CFU enumeration were collected under sterile conditions at the time of inoculation and at the time of approximately maximum current density. Samples were serially diluted (1:1000 to 1:1000000) in sterile PBS, and plated on MRS for CFUs enumeration after overnight incubation at 30 °C. Dry weight was determined using a 25 mL sample collected at approximately maximum current density. Cells were harvested by centrifugation (5250 g, 12 min, 4 °C) and washed twice in 50 mL ddH_2_O. Afterwards cells were resuspended in 1mL of ddH_2_O and transferred to microfuge tubes (previously weighted). Cells were harvested by centrifugation (5250 g, 12 min, 4 °C), and the tubes were then transferred to an evaporator to remove humidity. The microfuge tubes were then cooled in a desiccator for 30 min and the weight of each tube was measured to determine cell weight. The difference between the weight of each tube with the pellet and before containing it allowed us to determine the dry weight/mL.

### RNA-seq library construction and transcriptome analysis

*L. plantarum* NCIMB8826 was grown in triplicate to exponential phase (OD_600_ 1.0) at 37 °C in mMRS with or without the supplementation of 20 µg/mL DHNA and 1.25 mM ferric ammonium citrate. Cells were collected by centrifugation at 10,000 g for 3 min at 4 °C, flash frozen in liquid N_2_ and stored at −80 °C prior to RNA extraction as previously described (Golomb et al., 2016). Briefly, frozen cell pellets were resuspended in cold acidic phenol:chloroform:isoamyl alcohol (pH 4.5) [125:24:1] (Invitrogen, Carlsbad, CA, USA) before transferring to 2 mL screw cap tubes containing buffer (200 mM NaCl, 20 mM EDTA), 20% SDS, and 300 mg 0.1 mm zirconia/silica beads. RNA was extracted by mechanical lysis with an MP Fastprep bead beater (MP Biomedicals, Santa Ana, CA, USA) at 6.5 m/s for 1 min. The tubes were centrifuged at 20,000 g at 4 °C for 3 min and the upper aqueous phase was transferred to a new tube. The aqueous phase was extracted twice with chloroform:isoamyl alcohol [24:1] (Fisher Scientific, Waltham, MA, USA), The aqueous phase was then transferred to a new tube for RNA ethanol precipitation (Green and Sambrook, 2020). RNA was then quantified on a Nanodrop 2000c (ThermoFisher), followed by double DNAse digestion with the Turbo DNA-free Kit (Invitrogen) according to the manufacturer’s protocols. The quality of the remaining RNA was checked using a Bioanalyzer RNA 6000 Nano Kit (Agilent Technologies, Santa Clara, CA, USA) (all RIN values > 9) and then quantified with the Qubit 2.0 RNA HS Assay (Life Technologies, Carlsbad, CA, USA). For reverse-transcription PCR (RT-PCR), 800 ng RNA was converted to cDNA with the High Capacity cDNA Reverse Transcription Kit (Applied Biosystems, Foster City, CA, USA) according to the manufacturer’s protocols. Quantitative RT-PCR was performed on a 7500 Fast Real-Time PCR System (Applied Biosystems) using the PowerUp SYBR Green Master Mix (ThermoFisher) and RT-PCR primers listed in **Table S3**. The 2-ΔΔCt method was used for relative transcript quantification using *rpoB* as a control (Livak and Schmittgen, 2001).

For sequencing, ribosomal-RNA (rRNA) was depleted from 4 µg RNA using the RiboMinus Eukaryote Kit v2 with specific probes for prokaryotic rRNA (ThermoFisher) following the manufacturer’s instructions. RNA was then fragmented to approximately 200 bp, converted to cDNA, and barcoded using the NEBnext Ultra-directional RNA Library Kit for Illumina (New England Biolabs, Ipswitch, MA, USA) with NEBnext Multiplex Oligos for Illumina (Primer Set 1) (New England Biolabs) following the manufacturer’s protocols. cDNA libraries containing pooled barcoded samples was run across two lanes of a HiSeq400 (Illumina, San Diego, CA, USA) on two separate runs for 150 bp paired-end reads (http://dnatech.genomecenter.ucdavis.edu/). An average of 36,468,428 raw paired-end reads per sample was collected (**Table S4**). The DNA sequences were quality filtered for each of the 12 samples by first visualizing with FastQC (ver. 0.11.8) (Andrews and Others, 2010) to check for appropriate trimming lengths, followed by quality filtering with Trimmomatic (ver. 0.39) (Bolger et al., 2014). Remaining reads then were aligned to the NCIMB8826 chromosome and plasmids using Bowtie2 (ver. 2.3.5) in the [-sensitive] mode (Langmead and Salzberg, 2012). The resulting “.sam” files containing aligned reads from Bowtie2 were converted to “.bam” files with Samtools (ver 1.9) (Li et al., 2009) before counting aligned reads with FeatureCounts in the [-- stranded=reverse] mode (ver. 1.6.4) (Liao et al., 2014). Reads aligning to noncoding sequences (e.g., rRNA, tRNA, trRNA, etc.) were excluded for subsequent analyses. Differential gene expression based on culture condition was determined with DESeq2 (Love et al., 2014) using the Wald test in the R-studio shiny app DEBrowser (ver 1.14.2) (Kucukural et al., 2019).

Differential expression was considered significant with a False-discovery-rate (FDR)-adjusted *p-*value ≤ 0.05 and a Log_2_ (fold-change) ≥ 0.5. Clusters of Orthologous Groups (COGs) were assigned to genes based on matches from the eggNOG (ver. 5.0) database (Huerta-Cepas et al., 2019).

### Redox probe assays

Hamilton oxidation-reduction potential (ORP) probes (Hamilton Company, Reno, NV, USA) were inserted into air-tight Pyrex (Corning Inc., Corning, NY, USA) bottles containing mMRS supplemented with 20 µg/mL DHNA and 1.25 mM ferric ammonium citrate and incubated in a water bath at 37 °C. A custom cap for the Pyrex bottles was 3D printed with polylactic acid filament (2.85 mm diameter) such that the ORP probe threads into the cap and an o-ring seal can be used to provide an air-tight seal between the probe and the cap. The ORP was allowed to equilibrate over 40 min before *L. plantarum* NCIMB8826, MLES100, or MLES101 were inoculated at an OD600 of 0.1. Two uninoculated controls were used to measure baseline ORP over time. The ORP data was collected via Modbus TCP/IP protocol (Stride Modbus Gateway, AutomationDirect, Cumming, GA, USA) into a database (OSIsoft, San Leandro, CA, USA) and analyzed in MATLAB (Mathworks, Nantick, MA, USA). pH was measured using a Mettler Toledo SevenEasy pH meter (Mettler Toledo, Columbus, OH, USA). Cells were collected at either 24 h or at the greatest ORP difference between the wild-type and mutant strains (ΔmV_max_) by centrifugation at 10,000g for 3 min and used for ferrihydrite reduction analyses.

### ATP and NAD^+^/NADH quantification

Frozen cell pellets were suspended in PBS and lysed by mechanical agitation in a FastPrep 24 (MP Biomedicals) at 6.5 m/sec for 1 min. The cell lysates were then centrifuged at 20,000 g for 3 min at 4°C. ATP and NAD^+^ and NADH in the supernatants were then quantified with the Molecular Probes ATP Quantification Kit (ThermoFisher) and the Promega NAD/NADH-Glo Kit (Promega, Madison, WI, USA), respectively according to the manufacturers’ instructions.

### Kale juice fermentation assay

Green organic kale, purchased from a market (Whole Foods), was washed with tap water and air dried for 1 h as previously recommended (Kim, 2017-9). A total of 385 g of the leaves and stems were shredded with an electric food processor in 1 L ddH20. The kale juice was then diluted with 0.35 L ddH2O and autoclaved (121°C, 15 min). The juice was then centrifuged under sterile conditions at 8000 rpm for 20 min and the supernatant was collected. A rifampicin-resistant variant of *L. plantarum* NCIMB8826-R (Tachon et al., 2014) (grown for 19 h in MRS medium at 37 °C, 50 µg Rif/mL) was inoculated to an estimated final OD of approximately 0.05, and DHNA (20 µg/mL) was added where appropriate. Cells were collected and washed as previously described for the bioelectrochemical assays in mCDM. The anodic chambers of bioreactors assembled as previously described (anode of 4.3*6 cm) were filled with 125 mL of the inoculated kale juice and incubated at 30°C purged with N_2_. After 1 h, the anodes were polarized to 0.2 V versus Ag/AgCl (sat. KCl) (EET conditions) or kept at open circuit (OC, no EET). Viable cells were measured by plating 10-fold serial dilutions in MRS agar plates with 50 µg/mL of Rif.

### Calculations

The total electrons harvested on the anode were estimated by integrating the area (charge) under the chronoamperometric curve (current response (A) over time (s)), which was corrected by subtracting the current baseline obtained before *L. plantarum* was added to the system. This obtained charge was then converted to mol of electrons using the Faraday constant (96,485.3 A*s/mol electrons).

### Data accession numbers

*L. plantarum* RNA-seq data are available in the NCBI Sequence Read Archive (SRA) under BioProject accession no. PRJNA717240. A list of the completed Lactobacillales genomes used in the DNA sequence analysis is available in the Harvard Dataverse repository at https://doi.org/10.7910/DVN/IHKI0C.

## ACKNOWLEDGEMENTS

This work was supported by the National Science Foundation grant #1650042 and Office of Naval Research grant 0001418IP00037 (CMAF). Work at the Molecular Foundry was supported by the Office of Science, Office of Basic Energy Sciences, of the U.S. Department of Energy under Contract No. DE-AC02-05CH11231. James Nelson was supported by the Rodgers University fellowship in Electrical and Computer Engineering.

## ADDITIONAL INFORMATION

### Author details

Sara Tejedor-Sanz

Department of Biosciences, Rice University, Houston, United States and Lawrence Berkeley National Laboratory, Berkeley, United States Contribution: Conceptualization, Investigation, Data curation, Formal analysis, Validation, Visualization, Methodology, Writing - original draft preparation, review and editing Competing interests: No competing interests declared

Eric T. Stevens

Department of Food Science & Technology, University of California-Davis, Davis, United States Contribution: Conceptualization, Investigation, Data curation, Formal analysis, Validation, Visualization, Methodology, Writing - original draft preparation, review and editing Competing interests: No competing interests declared

Peter Finnegan

Department of Food Science & Technology, University of California-Davis, Davis, United States Contribution: Software, Data curation, Investigation, Writing - review and editing. Competing interests: No competing interests declared

James Nelson

Department of Electrical and Computer Engineering, University of California-Davis, Davis, United States Contribution: Software, Data curation, Investigation, Writing - review and editing Competing interests: No competing interests declared

Andre Knoessen

Department of Electrical and Computer Engineering, University of California-Davis, Davis, United States Contribution: Software, Data curation, Methodology, Resources, Writing - review and editing Competing interests: No competing interests declared

Samuel H. Light

Department of Microbiology, University of Chicago, United States Contribution: Conceptualization, Writing - review and editing Competing interests: No competing interests declared

Caroline M. Ajo-Franklin

Department of Biosciences, Rice University, Houston, United States and Lawrence Berkeley National Laboratory, Berkeley, United States Contribution: Conceptualization, Funding Acquisition, Supervision, Methodology, Writing - original draft preparation, Writing - review and editing Competing interests: No competing interests declared

Maria L. Marco

Department of Food Science & Technology, University of California-Davis, Davis, United States Contribution: Conceptualization, Funding Acquisition, Supervision, Methodology, Writing - original draft preparation, Writing - review and editing Competing interests: No competing interests declared

## Supporting Information

**Fig S1.**
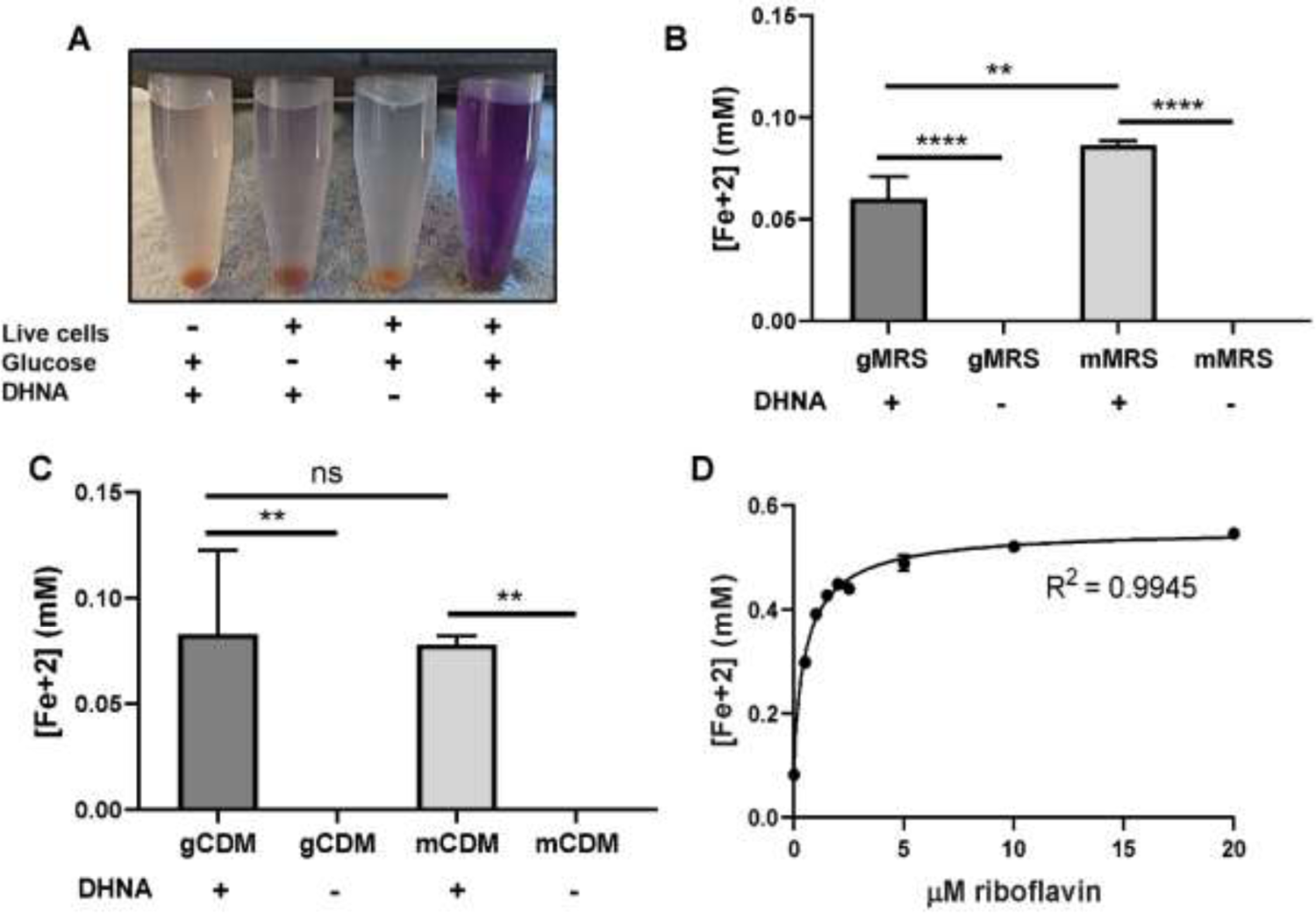
Iron reduction by *L. plantarum* is dependent upon DHNA, carbon source, and riboflavin. Reduction of Fe^3+^ (ferrihydrite) to Fe^2+^ by *L. plantarum* NCIMB8826 after growth in **(A and B)** glucose-containing MRS (gMRS) or **(C)** CDM (gCDM). Fe^3+^ reduction capacity was measured by the ferrihydrite reduction assay. **(A)** For inactivation, cells were incubated at 85 ^°^C in PBS for 30 min prior to the assay. Significant differences in iron reduction were determined by one-way ANOVA with Tukey’s post-hoc test (n = 3), ** p ≤ 0.01; *** p ≤ 0.001; **** p ≤ 0.0001. **(D)** Reduction of Fe^3+^ to Fe^2+^ by *L. plantarum* after 3 h in the presence of 20 μg/mL DHNA, 55 mM mannitol, and increasing concentrations of riboflavin. The average ± standard deviation of three replicates is shown.

**Fig S2.**
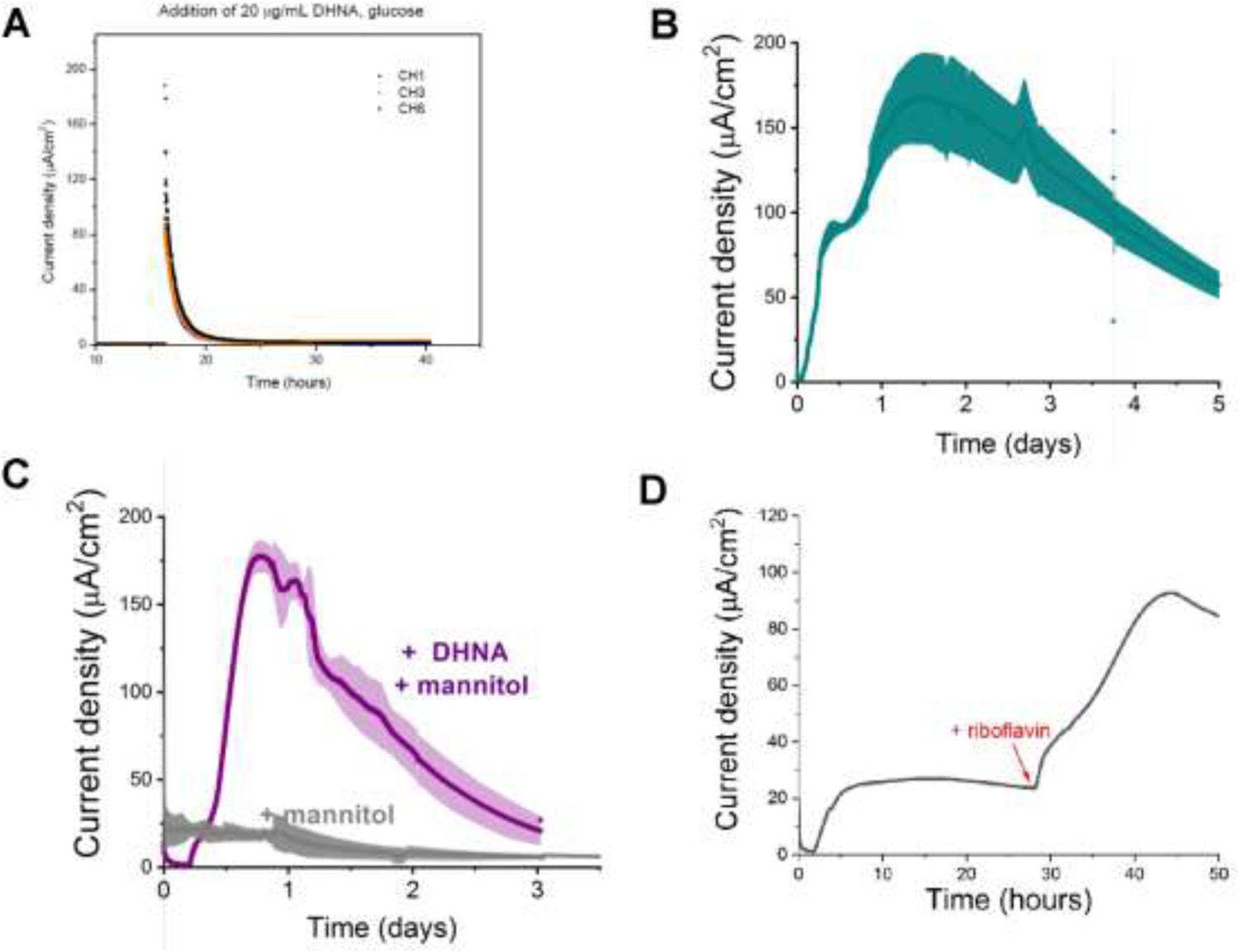
Current production by *L. plantarum* is a biotic process dependent on DHNA, carbon source, and riboflavin. **(A)** Abiotic current density response in bioelectrochemical reactors over time in mannitol-containing MRS (mMRS) upon DHNA (20 μg/mL) addition. **(B)** Current density produced by *L. plantarum* in mMRS with 20 μg/mL DHNA. **(C)** Current density produced by *L. plantarum* in gMRS with 20 μg/mL DHNA. **(D)** Effect of riboflavin addition on current density production by *L. plantarum* in mannitol-containing CDM (mCDM) in the presence of 20 μg/mL of DHNA. The average ± standard deviation of three replicates is shown.

**Fig S3.**
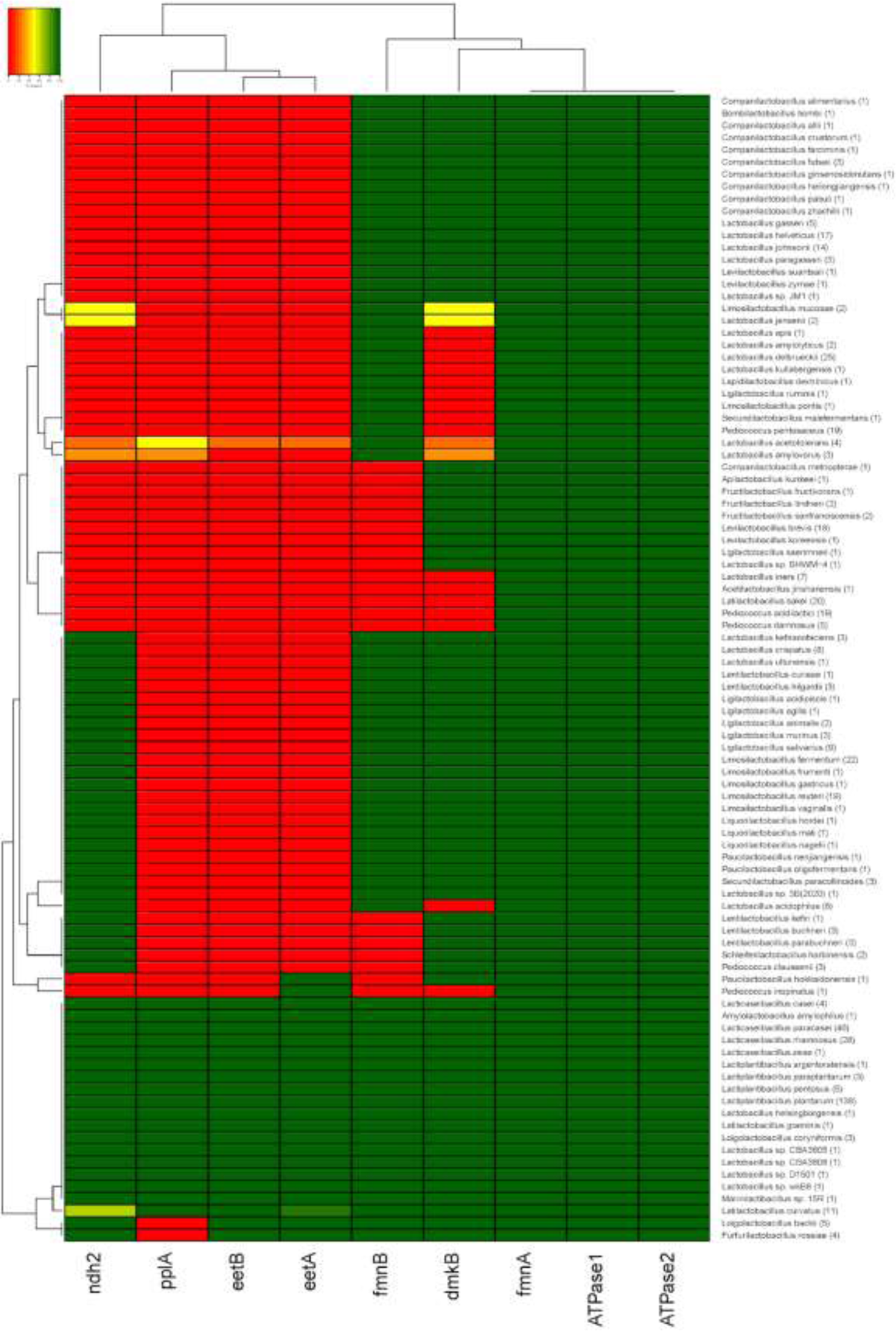
Conservation of FLEET locus genes among lactobacilli. Heatmap showing the percentage of bacteria in the *Lactobacillus*-genus complex containing genes in the FLEET locus. Homology searches were conducted using tBLASTx for 1,788 complete LAB genomes in NCBI (downloaded 02/25/2021) against the *L. plantarum* NCIMB8826 FLEET locus. A match was considered positive with a Bit-score ≥ 50 and an E-value of ≤ 10^-3^.

**Fig S4.**
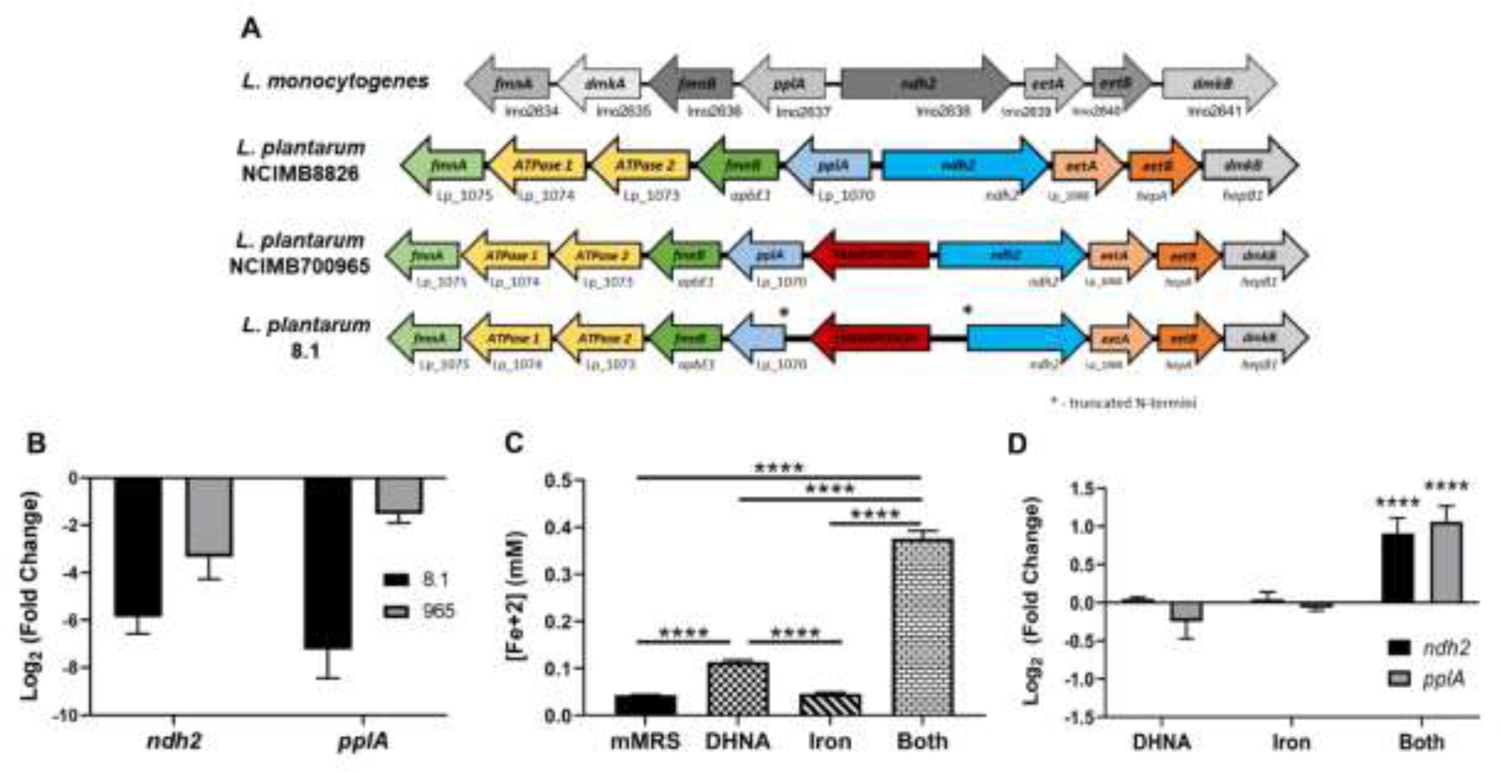
*ndh2* and *pplA* are required for iron reduction through EET. **(A)** Visualization of the FLEET locus in *L. monocytogenes* and three strains of *L. plantarum*. Genes are annotated based on predicted functions within the FLEET system. **(B)** Relative expression of *ndh2* and *pplA* in *L. plantarum* strains 8.1 and NCIMB700965 (“965”) during growth in MRS compared to *L. plantarum* NCIMB8826. The average ± standard deviation of three replicates is shown. **(C)** Reduction of Fe^3+^ (ferrihydrite) to Fe^2+^ by *L. plantarum* after growth to mid-exponential phase in mMRS with or without the supplementation of 20 μg/mL DHNA, 1.25 mM ferric ammonium citrate (“iron”), or both before being subject to the ferrihydrite reduction assay. **(D)** Relative expression of *ndh2* and *pplA* in *L. plantarum* during growth in mMRS containing 20 μg/mL DHNA, 1.25 ferric ammonium citrate (“iron”), or both compared to during growth in mMRS. Significant differences determined through **(C)** One-way ANOVA with Tukey’s post-hoc test (n = 3) or **(D)** Two-way ANOVA with Sidak’s post-hoc test (n = 3), **** p ≤ 0.0001.

**Fig S5.**
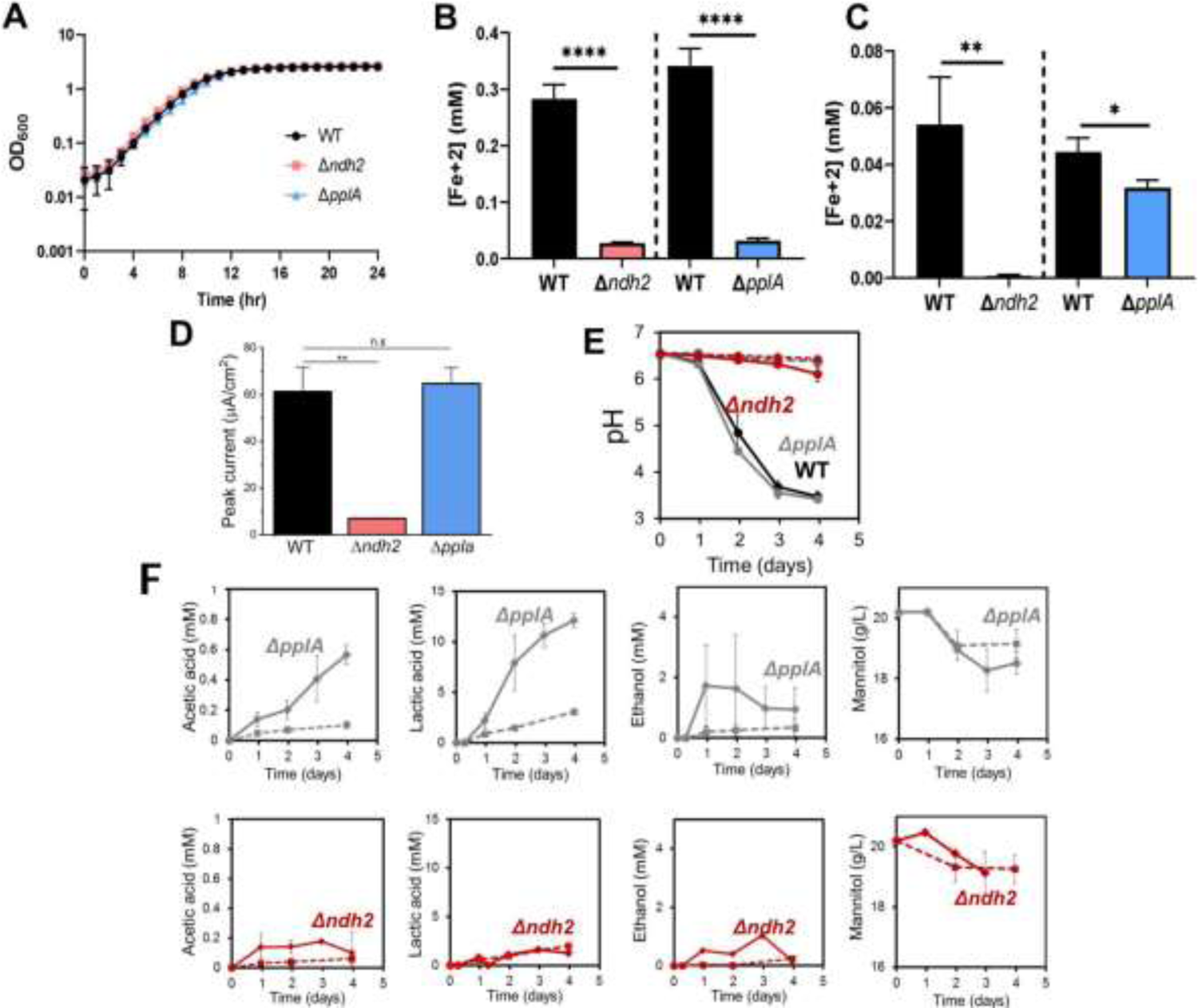
Impact of *ndh2* and *pplA* deletion on growth, iron reduction, current density and metabolites production. **(A)** Growth of wild-type *L. plantarum*, Δ*ndh2*, or Δ*pplA* in mMRS supplemented with 20 μg/mL DHNA and 1.25 mM ferric ammonium citrate. **(B and C)** Reduction of Fe^3+^ (ferrihydrite) to Fe^2+^ by *L. plantarum* or FLEET deletion mutants from Fig 3B in the ferrihydrite reduction assay at **(B)** ΔmVmax or **(C)** stationary phase. **(D)** Peak current generated by *L. plantarum* wild-type, Δ*ndh2*, and Δ*pplA* from Fig 3C. **(E)** pH measurements and **(F)** metabolites produced in the bioelectrochemical reactors inoculated with wild-type *L. plantarum,* Δ*ndh2*, or Δ*pplA* from Fig 3C. Solid lines denote the presence of an anode polarized to +0.2V (vs Ag/AgCl sat. KCl) while dashed lines denote open circuit conditions. The average ± standard deviation of three replicates is shown. Significant differences determined through **(B and C)** Two-tailed t-test (n = 3) or **(D)** One-way ANOVA with Dunn-Sidak post-hoc test (n = 3), * p ≤ 0.05; ** p ≤ 0.01; *** p ≤ 0.001; **** p ≤ 0.0001.

**Fig S6.**
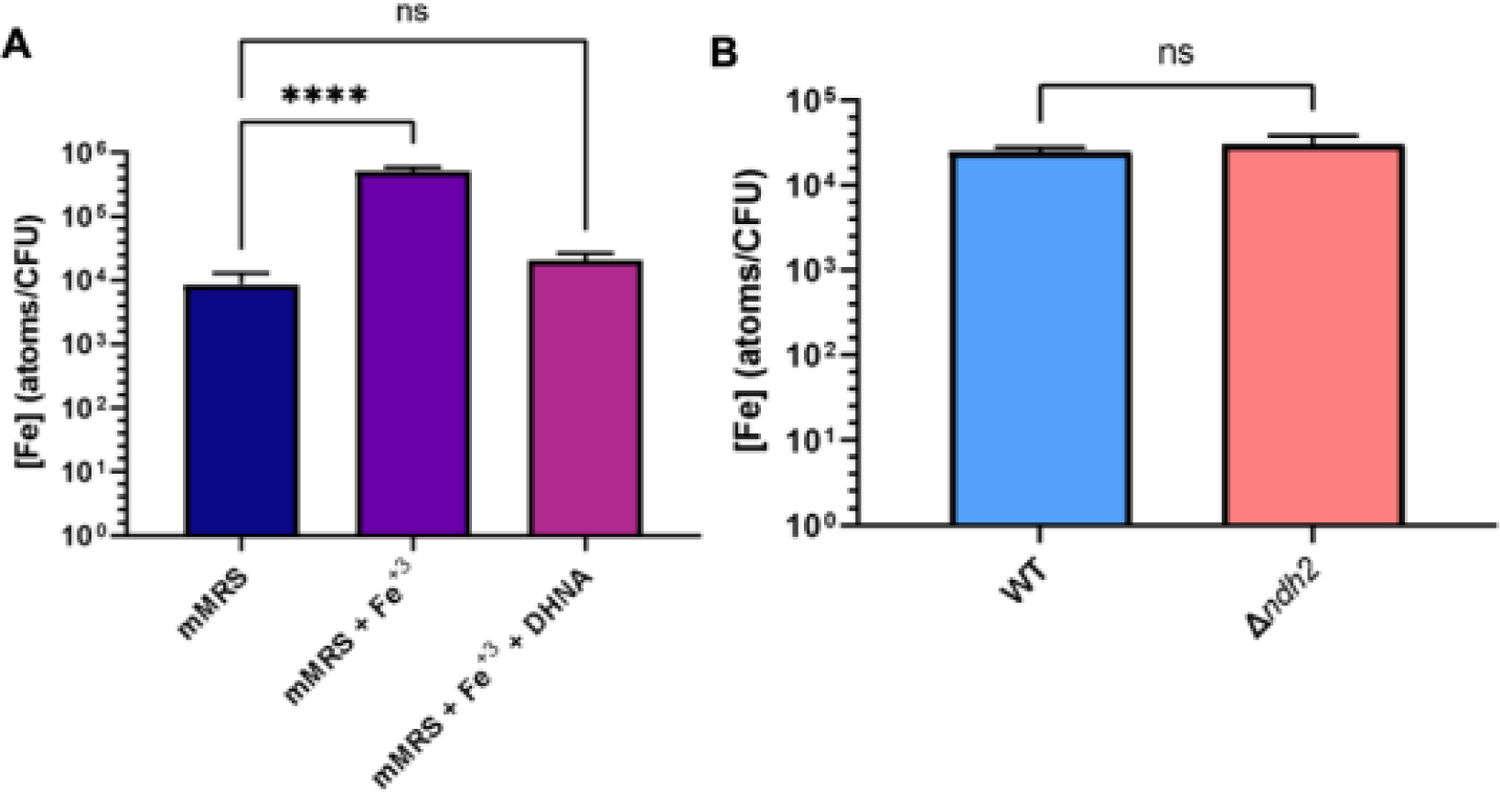
Intracellular iron concentrations in *L. plantarum* are not affected by EET. Inductively coupled plasma mass spectrometry (ICP-MS) quantification of intracellular iron from **(A)** *L. plantarum* grown 18 h in mMRS containing 1.25 mM ferric ammonium citrate (“Fe^+3^”) or 1.25 mM ferric ammonium citrate and 20 μg/mL DHNA. **(B)** Wild-type *L. plantarum* and *L. plantarum* Δ*ndh2* grown in mMRS supplemented with 1.25 mM ferric ammonium citrate and 20 μg/mL DHNA. Significant differences determined by **(A)** One-way ANOVA with Dunnett’s post-hoc test (n = 3) or **(B)** Two-tailed t-test, **** p ≤ 0.0001.

**Fig S7.**
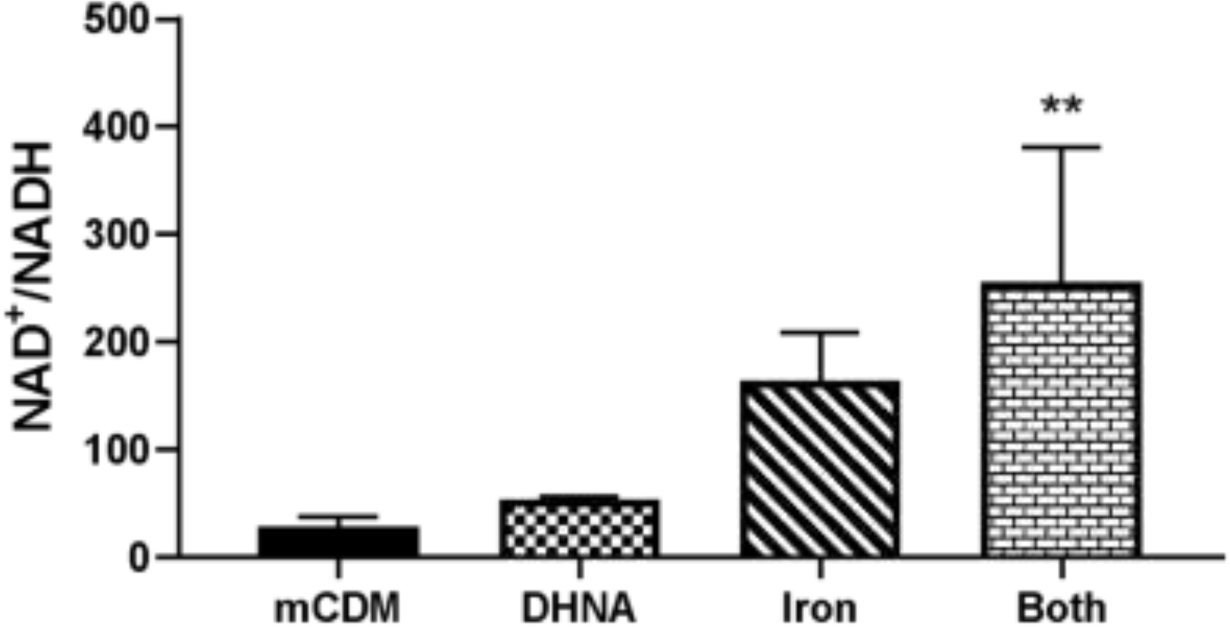
Use of Fe^3+^ as an electron acceptor allows *L. plantarum* to regenerate NAD^+^. NAD^+^/NADH ratios of *L. plantarum* grown to mid-exponential phase in mCDM with/without the supplementation of 20 μg/mL DHNA, 1.25 mM ferric ammonium citrate (“iron”), or both components. Significant differences determined through One-way ANOVA with Dunnett’s post-hoc test (n = 3), ** p ≤ 0.01.

**Fig S8.**
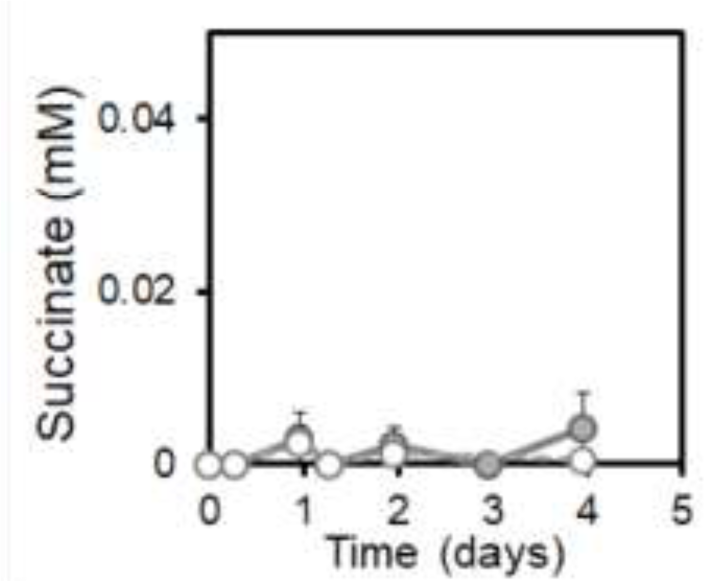
EET by L. plantarum does not involve TCA cycle metabolites. **(A)** Succinate production under EET (solid line) and open-circuit (dashed line) conditions from Fig 5. The average ± standard deviation of three replicates is shown.

**Fig S9.**
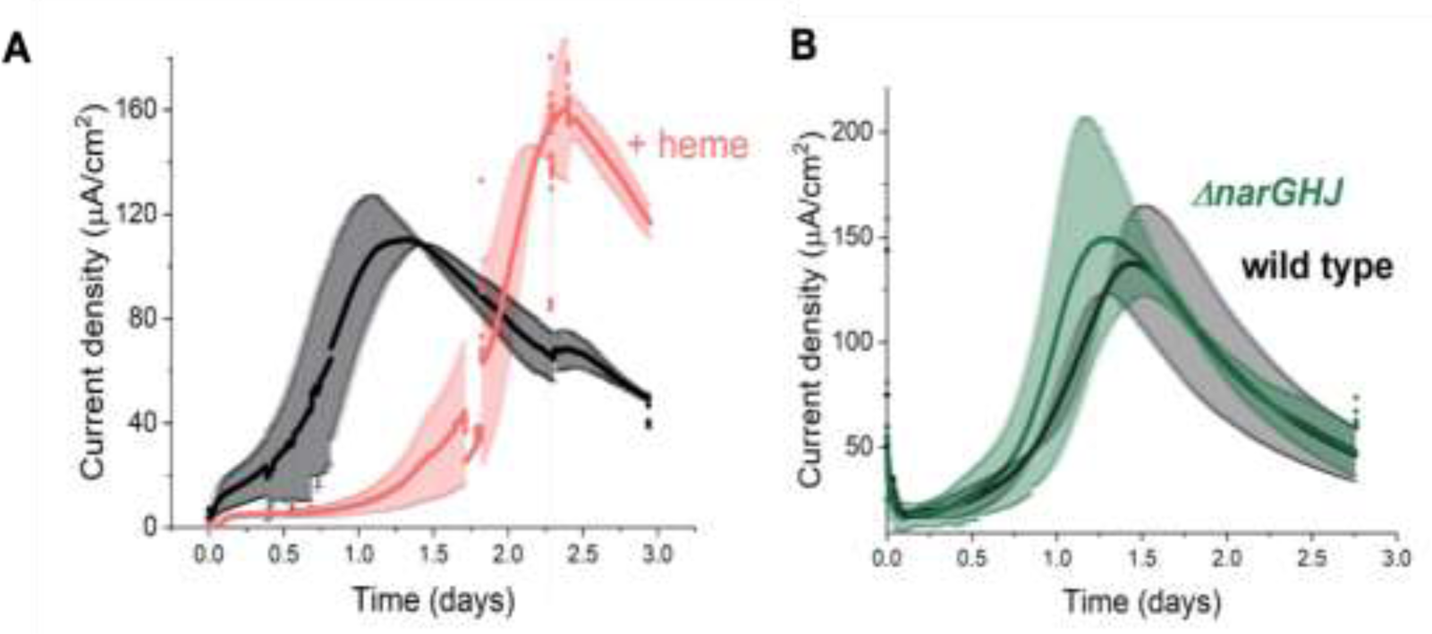
EET by *L. plantarum* is not dependent on aerobic or anaerobic respiration components. **(A)** Effect on *L. plantarum* current production with heme addition to reconstitute the aerobic electron transport chain. **(B)** Effect on *L. plantarum* current production with the deletion of the nitrate reductase A. The anode was polarized to +0.2V (vs Ag/AgCl sat. KCl). The average ± standard deviation of two replicates is shown.

**Fig S10.**
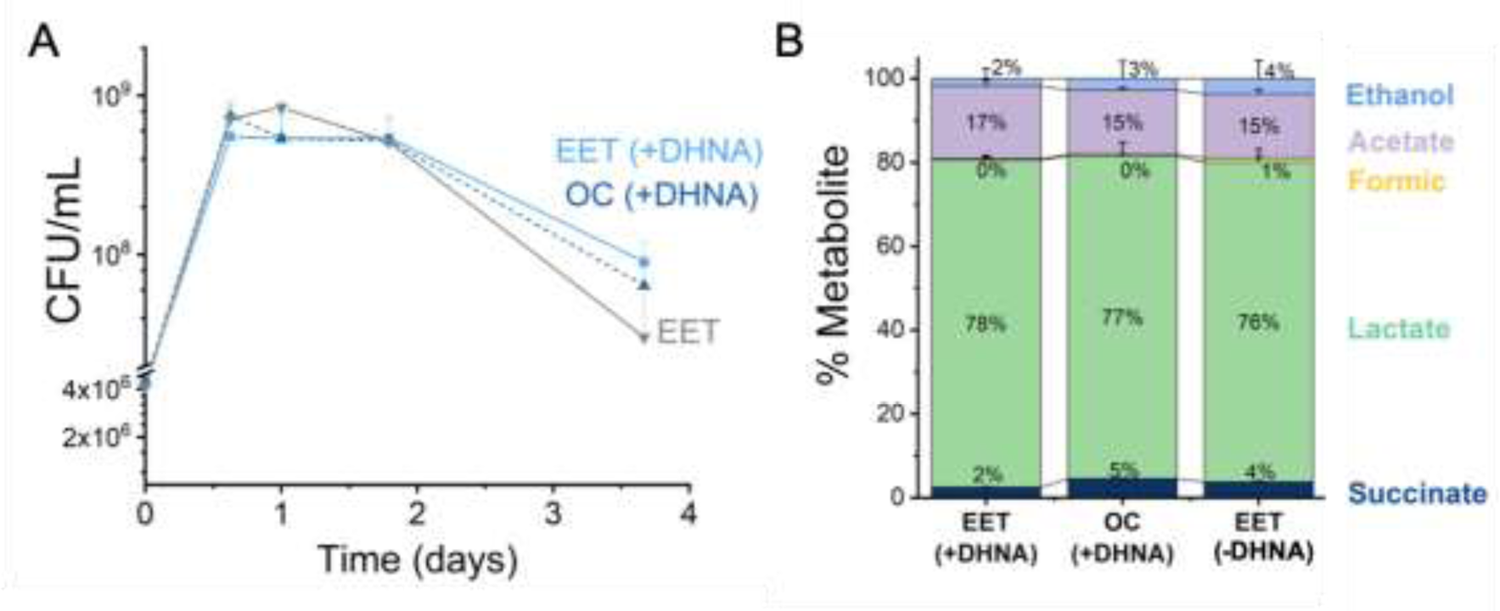
EET does not impact cell viability and distribution of metabolites in a kale fermentation. **(A)** Viable cells of *L. plantarum* NCIMB8826-R during the fermentation of kale juice in the presence of a polarized anode with/without DHNA, and under open circuit conditions with DHNA. **(B)** Distribution of metabolites after 2 days of kale juice fermentation. The anode polarization was maintained at +0.2 V (vs Ag/AgCl sat. KCl). The average ± standard deviation of three replicates is shown.

## Supporting Tables

**Table S1.** **Strains and plasmids used in this study.**

| Strains | Isolation Source / Description | Reference |
| --- | --- | --- |
| *L. plantarum* NCIMB8826 | Human saliva | (Dandekar, 2019) |
| *L. plantarum* NCIMB8826-R | Rifampicin-resistant mutant of NCIMB8826 | (Yin et al., 2018) |
| *L. plantarum* MLES100 | Deletion mutant of NCIMB8826 lacking *ndh2* | This study |
| *L. plantarum* MLES101 | Deletion mutant of NCIMB8826 lacking *pplA* | This study |
| *L. plantarum* MLEY100 | Deletion mutant of NCIMB8826 lacking *narGHIJ* | This study |
| *L. plantarum* B1.3 | Brown flour teff injera | (Yu et al., 2021) |
| *L. plantarum* AJ11 | Fermented olives; commercial fermentation | (Yu et al., 2021) |
| *L. plantarum* 8.1 | Wheat boza | (Yu et al., 2021) |
| *L. plantarum* ATCC 202195 | Human stool | (Wright et al., 2020) |
| *L. plantarum* NCIMB700965 | Cheese | (Heeney and Marco, 2019) |
| *L. pentosus* BGM48 | Fermented olives | (Golomb et al., 2013) |
| *L. casei* BL23 | Dairy products | (Mazé et al., 2010) |
| *L. brevis* ATCC 367 | Silage | (Makarova et al., 2006) |
| *L. lactis* KF147 | Mung bean sprouts | (Siezen et al., 2010) |
| *L. lactis* IL1403 | Cheese | (Bolotin et al., 2001) |
| *L. rhamnosus* GG | Human intestine | (Kankainen et al., 2009) |
| *L. murinus* ASF361 | Mice | (Wannemuehler et al., 2014) |
| *E. faecalis* ATCC 29212 | Urine | (Minogue et al., 2014) |
| *E. faecium* ATCC 8459 | Cheese | (Kopit et al., 2014) |
| *P. pentosaceus* ATCC 25745 | Plants | (Makarova et al., 2006) |
| *S. agalactiae* ATCC 27956 | Bovine udder infection | (McDonald and McDonald, 1976) |
| *E. coli* DH5α | *fhuA2 lac(del)U169 phoA glnV44 Φ80' lacZ(del) M15 gyrA96 recA1 relA1 endA1 thi-1 hsdR17*, amplification of cloning vector | (Taylor et al., 1993) |
| Plasmids |  |  |
| pRV300 | EryR AmpR, *E. coli* Ori pMB1, integrative vector | (Leloup et al., 1997) |
| pRV300:ndh2 | pRV300 derivative used for deletion of *ndh2* | This study |
| pRV300:pplA | pRV300 derivative used for deletion of *pplA* | This study |
| pRV300:narG | pRV300 derivative used for deletion of *narG* | This study |

**Table S2.** Chemically defined medium.

| **CDM component <sup>a</sup>** | **Final concentration (g/L)** |
| --- | --- |
| **Buffers and salts** |  |
| MOPS (3-(N-morpholino)propanesulfonic acid) | 8.371 |
| K<sub>2</sub>HPO<sub>4</sub> | 0.871 |
| NH<sub>4</sub>Cl | 1.070 |
| Na<sub>2</sub>SO<sub>4</sub> | 1.420 |
| **Metals** |  |
| MgCl<sub>2</sub> * 6H<sub>2</sub>O | 0.203 |
| MnCl<sub>2</sub> * 4H<sub>2</sub>O | 0.001 |
| FeSO<sub>4</sub> * 7H<sub>2</sub>O | 0.014 |
| **Amino acids** |  |
| Casamino acids | 3.000 |
| Cysteine-HCl * H<sub>2</sub>O | 0.145 |
| Tryptophan | 0.050 |
| **Wolfe's Vitamins<sup>b</sup>** |  |
| Pyridoxine HCl | 0.002 |
| Thiamine HCl | 0.001 |
| Riboflavin | 0.001 |
| Nicotinic acid | 0.001 |
| Calcium D-(+)-pantothenate | 0.001 |
| *p*-Aminobenzoic acid | 0.001 |
| Thioctic acid (α-Lipoic acid) | 0.001 |
| Biotin | 0.0004 |
| Folic acid | 0.0004 |
| Vitamin B12 | 0.00002 |
| **Wolfe's Minerals<sup>c</sup>** |  |
| Nitrilotriacetic acid (NTA) | 0.3 |
| MgSO<sub>4</sub> * 7H<sub>2</sub>O | 0.6 |
| MnSO<sub>4</sub> * H<sub>2</sub>O | 0.1 |
| NaCl | 0.2 |
| FeSO<sub>4</sub> * 7H<sub>2</sub>O | 0.02 |
| CoCl<sub>2</sub> * 6H<sub>2</sub>O | 0.02 |
| CaCl<sub>2</sub> | 0.02 |
| ZnSO<sub>4</sub> * 7H<sub>2</sub>O | 0.02 |
| CuSO<sub>4</sub> * 5H<sub>2</sub>O | 0.002 |
| AlK(SO)<sub>4</sub> * 12H<sub>2</sub>O | 0.002 |
| H<sub>2</sub>BO<sub>3</sub> | 0.002 |
| Na<sub>2</sub>MoO<sub>4</sub> * 2H<sub>2</sub>O | 0.002 |

<sup>a</sup> All solutions were prepared separately and sterile filtered through a 0.22 $\mu$ m filter before combining. Glucose or mannitol was also supplemented at 22.520 g/L or 22.772 g/L, respectively.
<sup>b</sup> pH adjusted to 11 before sterile filtering.
<sup>c</sup> After NTA addition, pH adjusted to 8 before adding remaining components and sterile filtering.

**Table S3.**
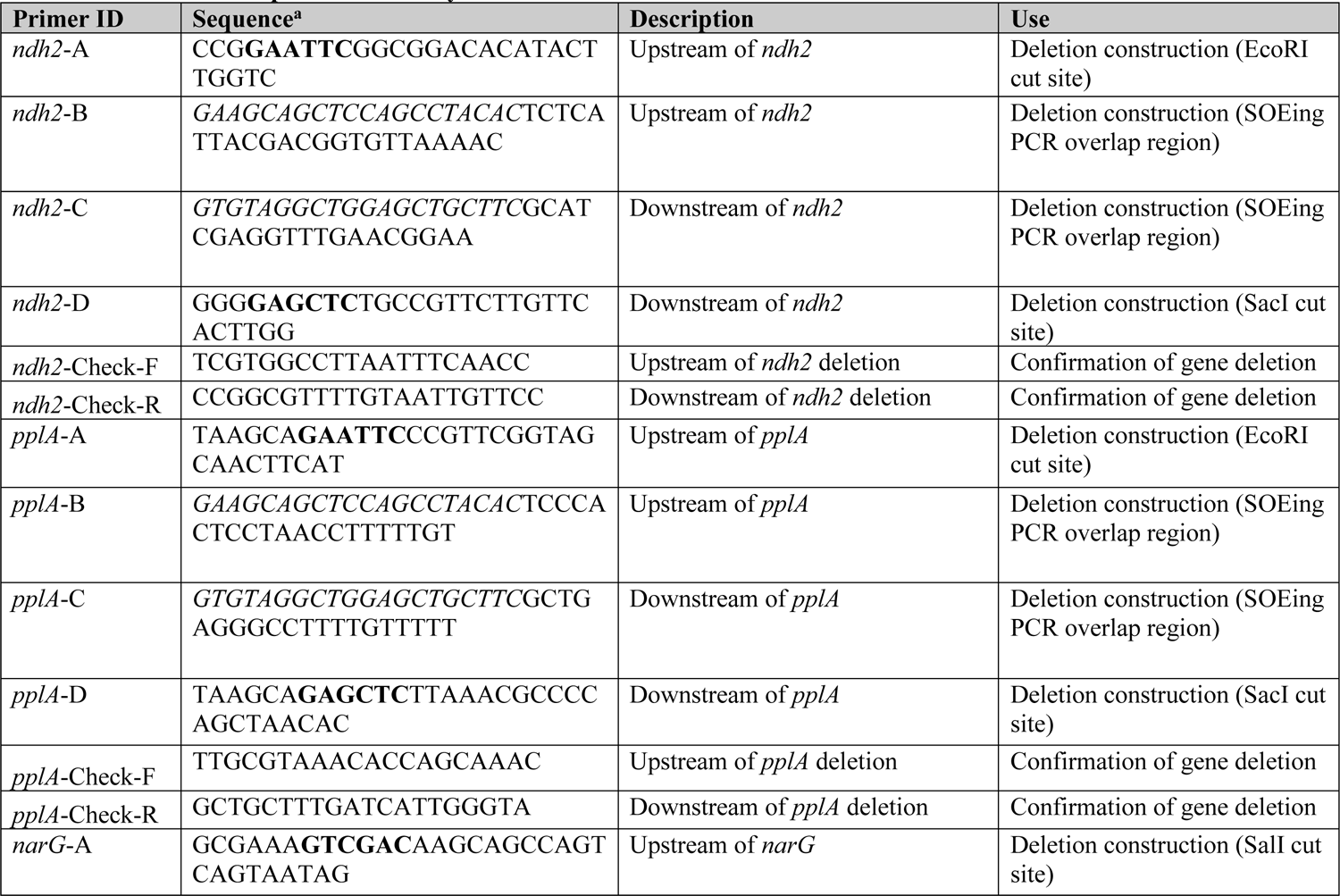
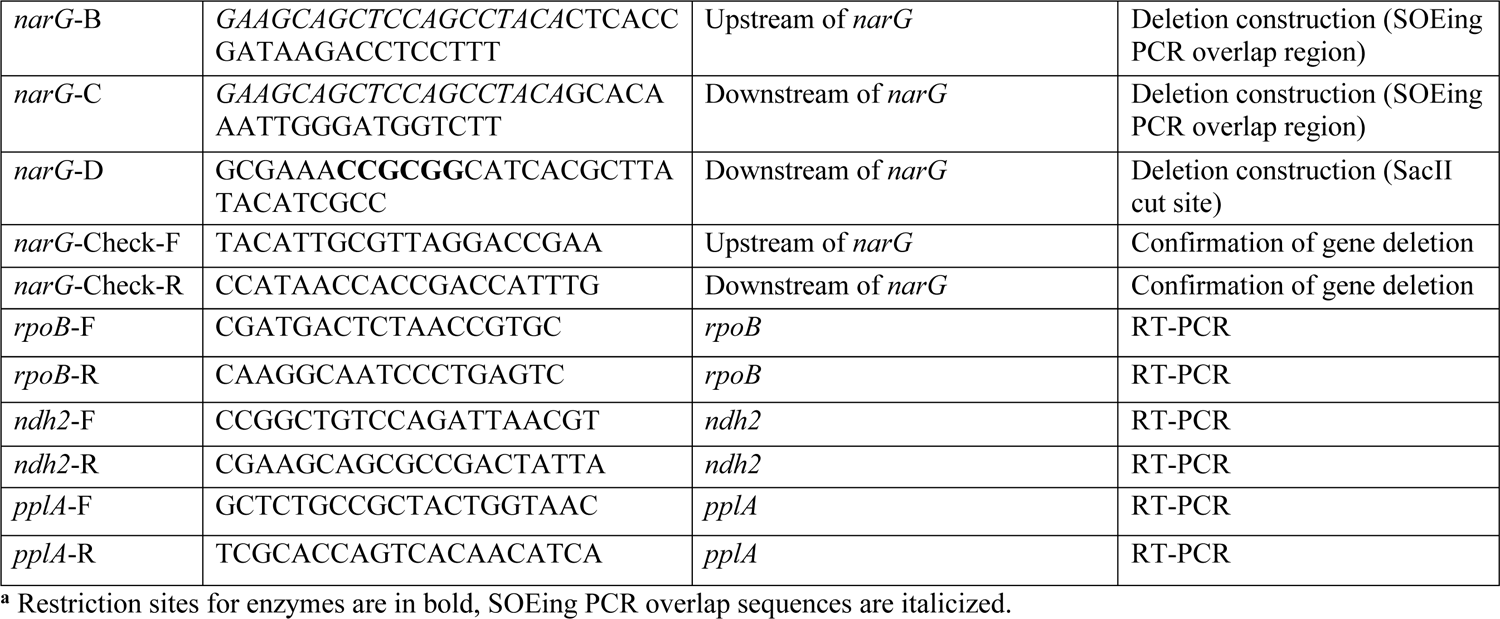
Primers developed for this study.

**Table S4.** Transcriptome read counts, alignment rate, and gene assignment rate.

| Sample ID | Total paired reads | Paired reads passed quality filtering<sup>a</sup> | Reads passed quality control (%) | Reads aligned to *Lp*<sup>b</sup> | Alignment (%) | Assigned reads<sup>c</sup> | Read assignment rate (%) | rRNA/tRNA/ncRNA Reads | Remaining reads for DE<sup>d</sup> | Remaining reads for DE<sup>d</sup> (%) |
| --- | --- | --- | --- | --- | --- | --- | --- | --- | --- | --- |
| mMRS-1 | 3.91E+07 | 3.40E+07 | 86.8 | 3.38E+07 | 99.6 | 30287036 | 89.6 | 10670771 | 19616265 | 64.8 |
| mMRS-2 | 3.96E+07 | 3.46E+07 | 87.2 | 3.44E+07 | 99.5 | 29762763 | 86.6 | 11736601 | 18026162 | 60.6 |
| mMRS-3 | 3.43E+07 | 3.03E+07 | 88.4 | 3.02E+07 | 99.5 | 26518101 | 87.8 | 13822383 | 12695718 | 47.9 |
| DHNA-1 | 4.13E+07 | 3.59E+07 | 87.0 | 3.57E+07 | 99.5 | 31800634 | 89.0 | 11364096 | 20436538 | 64.3 |
| DHNA-2 | 3.83E+07 | 3.35E+07 | 87.5 | 3.33E+07 | 99.5 | 29672126 | 89.0 | 9933102 | 19739024 | 66.5 |
| DHNA-3 | 3.99E+07 | 3.49E+07 | 87.5 | 3.47E+07 | 99.6 | 30954603 | 89.1 | 14806368 | 16148235 | 52.2 |
| Iron-1 | 3.31E+07 | 2.88E+07 | 86.8 | 2.86E+07 | 99.4 | 25147375 | 87.9 | 7180253 | 17967122 | 71.4 |
| Iron-2 | 2.47E+07 | 2.18E+07 | 88.2 | 2.16E+07 | 99.4 | 18943484 | 87.5 | 4976026 | 13967458 | 73.7 |
| Iron-3 | 3.10E+07 | 2.67E+07 | 86.1 | 2.66E+07 | 99.7 | 23221100 | 87.4 | 7577947 | 15643153 | 67.4 |
| Both-1 | 3.66E+07 | 3.03E+07 | 82.6 | 3.01E+07 | 99.6 | 26811334 | 88.9 | 12603853 | 14207481 | 53.0 |
| Both-2 | 4.39E+07 | 3.86E+07 | 87.9 | 3.85E+07 | 99.7 | 34163533 | 88.8 | 16841230 | 17322303 | 50.7 |
| Both-3 | 3.58E+07 | 3.11E+07 | 86.9 | 3.10E+07 | 99.7 | 27593814 | 89.0 | 14617870 | 12975944 | 47.0 |

<sup>a</sup> Paired-end reads filtered through Trimmomatic (ver. 0.39)
<sup>b</sup> Paired-end reads aligned to *L. plantarum* genome with Bowtie2 (ver. 2.3.5)
<sup>c</sup> Reads assigned to *L. plantarum* genes with FeatureCounts (ver. 1.6.4)
<sup>d</sup> Differential expression analyses done through DEseq2 on DEBrowser (ver. 1.14.2)

**Table S5.** Data used for calculating the bioenergetic balances.

|  | **Mannitol**<br>**consumed** | **Lactic acid** | **Acetic acid** | **Ethanol** | **Dry weight** | **Electrons**<br>**to anode** |
| --- | --- | --- | --- | --- | --- | --- |
|  | mM | mM | mM | mM | g/L | mM |
| EET<br>(anode) | 10.51 ± 3.14 | 6.32 ± 1.36 | 0.25 ± 0.03 | 0.47 ± 0.54 | 0.019 ± 0.04 | 7.04 ± 1.09 |
| OCP | 5.04±0.63 | 0.56 ± 0.01 | 0.035 ± 0.013 | 0.53 | 0.003 ± 0.001 | - |

**Table S6.** Comparison of the energy metabolism discovered in this study with fermentation in LAB and anaerobic respiration in Geobacter spp.

|  | **Homofermentation in LAB** | **Respiration in LAB** | **Anaerobic EET respiration in *Geobacter* spp.** | **Hybrid metabolism in LAB (this study)** |
| --- | --- | --- | --- | --- |
| Reduction of insoluble extracellular electron acceptor | No | No | Yes<br>4-8 mA/mg protein at peak (Marsili et al., 2010; Rose and Regan, 2015) | Both - soluble and insoluble<br>2 mA/mg- protein (at peak<sup>a</sup>) |
| NADH:quinone oxidoreductase is required for reduction of electron acceptor | No (Tachon et al., 2010) | Ndh1 (R. Brooijmans et al., 2009b, 2009b) | Yes<br>Ndh1 (proton pumping) (Chan et al., 2017) | Yes<br>Ndh2 (non-proton pumping) |
| Electron acceptor transcriptionally upregulates NADH dehydrogenase | No (this study) | No [aerobic respiration] (Pedersen et al., 2008) | Yes (Holmes et al., 2006) | Yes (with DHNA) |
| NAD<sup>+</sup>/NADH ratio | ~1 for glucose (Guo et al., 2017, p. 201)<br>~5 for mannitol (this study) | Near-zero for lactose (Johanson et al., 2020) | 10 (Fe<sup>3+</sup>)<br>10-100 (anode) (Rose and Regan, 2015; Song et al., 2016) | 260 (Fe<sup>3+</sup>)<br>45 (anode) |
| Fraction of electrons on electron acceptor | 40-98% pyruvate (Dirar and Collins, 1972) | 33% (nitrate) (R. J. W. Brooijmans et al., 2009, p. 1) | 75-90% anode (Speers and Reguera, 2012) | 20% anode<br>48% pyruvate |
| Metabolic flux is primarily through | Fermentative pathway (Bintsis, 2018) | Mixed-acid fermentation (R. Brooijmans et al., 2009a) | TCA cycle (Galushko and Schink, 2000; Mahadevan et al., 2006) | Fermentative pathway<br>- Mixed acid |
| ATP yield per substrate | 2~2-3 mol ATP/mol glucose for homolactic (Dirar and Collins, 1972)<br>0.75 mol ATP/mol mannitol (this study) | 3.3-3.9 mol ATP/mol lactose (Johanson et al., 2020) | ~9 mol ATP/mol substrate<sup>b</sup> (Mahadevan et al., 2006) | 1.5 mol ATP/mol mannitol |

<sup>a</sup> Calculated assuming that 50% of the dry cell weight is protein.
<sup>b</sup> *G. sulfurreducens* uses acetate as its electron donor. Since acetate is a 2 carbon electron donor, ATP yield is expressed per mol of a 6 carbon substrate.

## Notes

https://dataverse.harvard.edu/dataset.xhtml?persistentId=doi:10.7910/DVN/IHKI0C

### Competing Interest Statement

The authors have declared no competing interest.

